# Boosting of CAR-T cells with rhabdovirus is limited by type I interferon and rapid contraction

**DOI:** 10.1101/2024.11.29.626103

**Authors:** Rebecca Burchett, Claire G. Morris, Mira Ishak, Derek Cummings, Christopher L. Baker, Ricardo Marius, Natasha Kazdhan, Christopher M. Silvestri, John C. Bell, Brian D. Lichty, Scott R. Walsh, Yonghong Wan, Joanne A. Hammill, Jonathan L. Bramson

**Affiliations:** Centre for Discovery in Cancer Research, McMaster University, Hamilton, ON, Canada; McMaster Immunology Research Centre, McMaster University, Hamilton, ON, Canada; Department of Medicine, McMaster University, Hamilton, ON, Canada; Michael G. DeGroot Institute for Infectious Disease Research, McMaster University, Hamilton, ON, Canada; Ottawa Hospital Research Institute, Ottawa, ON, Canada

**Keywords:** Chimeric antigen receptor, oncolytic virus, cancer vaccines, adoptive T cell therapy

## Abstract

Rhabdovirus vaccines that encode tumour-associated antigens are potent boosting agents for adoptively transferred tumour-specific T cells. Employing rhabdovirus vaccines to boost adoptively transferred T cells relies on *a priori* knowledge of tumour epitopes, isolation of matched epitope-specific T cells, and a personalized vaccine, which limit clinical feasibility. Here, we investigated a universal strategy for boosting transferred tumour-specific T cells where boosting is provided through a chimeric antigen receptor (CAR) that is paired with a vesicular stomatitis virus (VSV) vaccine encoding the CAR-target. Boosting CAR-engineered tumour-specific T cells with paired VSV vaccines was associated with robust T cell expansion and delayed tumour progression in syngeneic models. CAR-T cell expansion and anti-tumour function was enhanced by blocking IFNAR1. However, vaccine-boosted CAR-T cells rapidly contracted and antigen-positive tumours re-emerged. In contrast, when the same T cells were boosted with VSV encoding antigen that stimulates through the TCR, the adoptively transferred T cells displayed improved persistence, tumour-specific endogenous cells expanded in parallel, and tumour cells carrying the antigen target were completely eradicated. Our findings underscore the need for further research into CAR-mediated boosting, how this differs mechanistically from TCR-mediated boosting, and the importance of engaging endogenous tumour-reactive T cells to achieve long-term tumour control.

## Introduction

Adoptive T cell therapy (ACT) has demonstrated remarkable clinical efficacy in the treatment of metastatic melanoma, HPV-induced cervical carcinoma, and hematological malignancies that are otherwise refractory to conventional therapies (1–7). Despite these successes, there remain biological and practical barriers to overcome, particularly in the treatment of resistant solid tumours where only modest clinical efficacy has been observed.

Successful ACT requires very large numbers of T cells, but spontaneously occurring tumour-specific T cells are rare. There are two main strategies for generating T cells for adoptive transfer. First is selection and *ex vivo* expansion of naturally occurring tumour-specific T cells that can be found in the peripheral blood and tumour tissues of cancer patients (8, 9). These tumour infiltrating lymphocytes (TIL) represent an endogenous T cell receptor (TCR) repertoire with specificity for multiple tumour targets. Large-scale *ex vivo* expansion of patient-derived TIL yields the large number of cells required for therapy; however, extended *ex* vivo culture often leads to exhaustion and terminal differentiation of the tumour-reactive T cells, thereby decreasing their antitumour potential (10). The second strategy functions by engineering a bulk T cell population to express a new tumour-specific TCR or a synthetic tumour-targeted receptor, such as a CAR (11). Large numbers of bulk T cells from peripheral blood can be engineered at once; however, this strategy typically yields a cell population with single antigen specificity. ACT targeting single antigens often fails to achieve long-term cures as a result of tumour heterogeneity and emergence of antigen loss variants (12). Therefore, a strategy that allows for robust expansion of functional multi-specific antitumour T cells is needed.

To address this, we have focused on strategies that exploit tools to promote *in vivo* expansion post-transfer. We have shown that oncolytic virus (OV) vaccines, wherein an OV is engineered to express a tumour-associated antigen, can be used to effectively boost tumour antigen-primed T cell populations *in vivo* (13–15). Similarly, when adoptive transfer of tumour-specific T cells is followed with an OV vaccine boost, we have observed robust *in vivo* expansion of the transferred cells and complete and durable tumour regressions in aggressive murine tumour models (16, 17). Neither the tumour-targeted ACT nor the OV vaccine alone was able to induce tumour regression in these models, demonstrating the therapeutic synergy of this approach.

Our previous studies have primarily focused on the evaluation of OV vaccines that boost T cells by engaging the native TCR, which is an MHC-restricted process that requires *a priori* knowledge of peptide epitopes. Clinical application of this approach would require that OV vaccines be personalized on a patient-by-patient basis - a process that is costly both in terms of time and resources. Here, we proposed an “off the shelf” approach to boosting where a universal OV vaccine is matched with a universal CAR that can be engineered into any T cell product. We hypothesized that tumour-specific T cells can be boosted through the CAR and achieve anti-tumour immunity via the TCR. Clinical experience has demonstrated that CAR T cells undergo robust expansion following ligation of their target antigen *in vivo* (18, 19). *In vivo* expansion of CAR-T cells is observed when ACT is combined with CAR-targeted lipid- or RNA-based vaccines, demonstrating that CARs can be successfully employed as boosting receptors (20–22). As such, CARs are well-positioned to mediate an OV vaccine boost.

Importantly, work from our lab, and others, has demonstrated that expression of a CAR does not impair native TCR function in engineered cells (23–26). The strategy we described herein exploits this dual functionality but takes a novel approach where the adoptively transferred T cells are boosted via the CAR and attack tumour cells via the TCR. We believe that this approach will allow for i*n vivo* expansion and differentiation of multi-specific populations of tumour-targeted T cells, such as TIL, in an antigen-agnostic manner.

Here, we have developed and validated a proof-of-concept syngeneic model to evaluate a CAR-mediated OV vaccine boost. We demonstrated that tumour-specific CD8+ T cells can be expanded in immunocompetent tumour-bearing recipients through engagement of a boosting CAR with a paired recombinant rhabdovirus vaccine. Mechanistic investigations revealed that the magnitude of the T cell expansion was limited by interferon-mediated suppression of viral replication. Even in the presence of interferon blockade, T cell expansion through the CAR was transient and tumours ultimately relapsed despite the continued expression of antigen. In contrast, boosting of the same T cells via their TCR promoted equally robust expansion but more durable persistence. While tumours ultimately relapsed in this setting, it was the result of antigen loss, which reinforces the improved therapeutic efficacy of T cells boosted through the TCR. Thus, while boosting through a CAR offers advantages with regards to pairing with a boosting vector, it may be disadvantaged by the non-canonical mechanisms of antigen presentation and T cell activation. Additionally, the simultaneous engagement of both endogenous and transferred tumour-reactive T cells with the booster vaccine appears to be critical for anti-tumour efficacy, something that is not achieved with a tumour-antigen agnostic “universal” CAR-boosting antigen.

## Materials and methods

### Sex as biological variable

Our study exclusively examined female mice. It is unknown whether the findings are relevant for male mice.

### Mice

C57BL/6 and BALB/c mice were purchased from Charles River Laboratories. IFNAR1 knockout (B6(Cg)-IFNAR1<tm1.2-Ees>/J) and C57BL/6 nude (B6.Cg-*Foxn1^nu^*/J) mice were purchased from The Jackson Laboratory. The P14-Thy1.1 strain was generated in-house by crossbreeding P14 mice (B6.Cg-*Tcra*^tm1Mom^ Tg(TcrLCMV)327Sdz; purchased from Taconic Laboratories, MMRRC stock #37394) onto a homozygous Thy1.1^+/+^ background (B6.PL-Thy1a/CyJ; purchased from Charles River Laboratories). DUC18 mice were kindly provided by Dr. Lyse Norian (University of Iowa).

### Cell lines

All cells were cultured at 37°C in a humidified atmosphere with 5% CO2. The human B cell leukemia line NALM-6, myeloma cell line KMS-11, and ovarian carcinoma cell line SKOV-3 were cultured in RPMI 1640 medium containing 8.7% FBS, 1.7 mM L-glutamine, 8.7 mM Hepes, 0.87x non-essential amino acids, 0.87 mM sodium pyruvate, 86.9 U/mL penicillin + 86.9 U/mL streptomycin, and 47.8 nM 2-mercaptoethanol. Platinum-E (PLAT-E) cells were maintained in DMEM supplemented with 8.9% FBS, 1.78 mM L-glutamine, 8.9 mM Hepes, 89 µg/mL Normocin, 8.9 µg/mL Blasticidin, and 0.89 µg/mL Puromycin. Parental CMS5 and CMS5 relapse (CMS5r) murine fibrosarcoma cell lines have been previously described (16, 27), CMS5r were used to generate a new cell line expressing the chimeric human BCMA-tNGFR antigen via lentiviral transduction (CMS5r-BCMA). CMS5, CMS5r-BCMA cells, MC38-gp33 murine colon adenocarcinoma cells expressing LCMV_GP33-41_ (KAVYNFATM) peptide (28), Vero cells, and HEK293T cells were maintained in Dulbecco’s Modified Eagle’s Media (DMEM) supplemented with 8.9% heat-inactivated fetal bovine serum (FBS), 0.89x GlutaMAX, 8.85 mM Hepes, and 88.5 U/mL penicillin + 88.5 µg/mL streptomycin. The murine melanoma line B16.F10-gp33 (29) was cultured in MEM/Earles medium supplemented with 8.6% FBS, 1x GlutaMAX, 8.61 mM Hepes, 86.1 U/mL penicillin + 86.1 µg/mL streptomycin, 0.86x MEM vitamin solution, 47.4 nM 2-mercaptoethanol, 0.86x non-essential amino acids, 0.86 nM sodium pyruvate, and 0.2 mg/mL G418 (Geneticin). B16.F10-gp33 NucLight Green (NLG) cells were generated from the parental line by transduction with commercially available lentivirus (Sartorius). Cell lines were regularly tested for mycoplasma contamination.

### Peptides and viral vectors

The H-2D^b^-restricted peptide of LCMV_GP33-41_ (KAVYNFATM) was purchased from GenScript Biotech and dissolved in PBS supplemented with 0.5% BSA. VSVΔ51 viruses encoding hBCMA-tNGFR (VSV-hBCMAtNGFR), hBCMA-mFc (VSV-hBCMAmFc), LCMV_GP33-41_ (VSV-gp33), Peptide H-2D^b^ Adpgk neoepitope (VSV-control), huBCMA-tNGFR_T2A_rsLuc (VSV-hBCMAtNGFR-rsLuc), and rsLuc_T2A_huBCMA-tNGFR (VSV-rsLuc-hBCMAtNGFR) were rescued in HEK293T cells, propagated in Vero cells, gradient purified, and titered as previously described (30).

### Generation of CAR retroviral vectors

The starting scaffold for all murine CAR vectors used in this study was generated in the Bramson lab and has been previously described (31). The two human CD19 targeted-CAR designs employed here differ in the linkers used for the ScFv. One uses a conventional glycine-serine linker and the other employs a Whitlow linker between the FMC63 V_H_ and V_L_ domains; the Whitlow linker includes a C-terminal proline residue that has been reported to increase stability and reduce aggregation of the ScFv (32). The described C11D5.3 ScFv was inserted into the murine CAR scaffold to generate human BCMA-targeted CARs (33). The murine 4-1BB CAR was generated by replacing the canonical TRAF-binding domains with the TRAF-binding domains from human 4-1BB, as previously described (34). Murine ecotropic gammaretrovirus was generated by transfection of Platinum-E (PLAT-E) packaging cells with 10 µg each of pRV2011 retroviral plasmid and pCL-Eco helper plasmid using Lipofectamine 2000 (Life Technologies), as previously described (35). Retroviral supernatants were collected at 48 hours post-transfection and concentrated 20x using Amicon Ultra 100K Centrifugal filters (EMD Millipore). Concentrated retroviral supernatants were aliquoted and stored at -80°C until use.

### Generation of murine CAR-T cells

Whole spleens were obtained from healthy 6-12 week-old Balb/c, C57BL/6, DUC18, or P14-Thy1.1 mice following isoflurane anaesthesia and euthanasia via cervical dislocation. Bulk splenocytes were released into T cell medium (RPMI 1640 (Gibco), 8.69% heat-inactivated fetal bovine serum (Gibco), 0.96x GlutaMAX, 8.69 mM HEPES (Roche), 0.87 mM sodium pyruvate (Sigma Aldrich), 0.87x non-essential amino acids (Gibco), 47.8 μM β-mercaptoethanol (Gibco), 86.9 U/mL penicillin + 86.9 μg/mL streptomycin (Gibco)) by gentle grinding of spleen tissue between glass microscope slides. Following erythrocyte lysis with ACK buffer, splenocytes were adjusted to a concentration of 3 x 10^6^ cells/mL in T cell medium containing 10 ng/mL rhIL-7 (Peprotech) and 2 µg/mL anti-mouse CD28 (clone 37.51, BioXCell) and were added to wells coated with anti-mouse CD3ε (clone 2C11, BioXCell) at 5 µg/mL. T cells were activated for 24 hours prior to retroviral transduction with CAR-retrovirus, as previously described (35). Cells were maintained at a concentration of 1-1.5 x 10^6^ cells/mL in medium containing 10 ng/mL rhIL-7 for the duration of *ex vivo* culture. 10 ng/mL rhIL-15 was included from day 4 post-activation to the end of culture. Cells were used in functional studies and cryopreserved for use in animal studies at 7-9 days post-activation.

### Flow cytometry antibodies and analysis

All antibodies used for flow cytometry are listed in **Table 1**. For murine T cells, splenocytes, and PBMCs, samples were first incubated for 15 minutes at 4°C in mouse BD Fc-block™ (rat anti-mouse CD16/CD32) at 2.5 µg/mL in FACS-EDTA buffer (PBS + 0.5% BSA + 2.5 mM EDTA). All surface antigen stains were prepared in FACS-EDTA buffer and incubated with cells at 4°C for 30-45 minutes. Recombinant human BCMA-Fc, recombinant human CD19-Fc, and recombinant human HER2-Fc (all R&D systems) were used to directly stain for BCMA-CAR, CD19-CAR, and HER2-CAR, respectively, followed by a fluorophore-conjugated secondary anti-human IgG. Viability staining was performed using the Molecular Probes LIVE/DEAD Fixable Near-IR dead cell stain kit (ThermoFisher Scientific). Cells were fixed and permeablized with BD Cytofix/Cytoperm prior to intracellular stains (Firefly Luciferase and Vaccinia B18R). Data were acquired on a BD LSRII (V/B/YG/R laser configuration), BD LSR Fortessa (V/B/R laser configuration), Beckman CytoFlex LX (NUV/V/YG/B/R laser configuration), or Beckman CytoFlex S (V/YG/B/R laser configuration) machine. Data were analysed using FlowJo (BD) and FCS Express 7 Research (De Novo Software Inc.) softwares.

### Intracellular cytokine staining

Murine T cells and PBMCs were stimulated with tumour targets or 1 µg/mL LCMV-GP_33– 41_ peptide (Biomer Technologies) in T cell media at 37°C for 5-6 hours. Brefeldin A (GolgiPlug, BD Biosciences; 1 μg/mL) was added for the last 4 hours of incubation. Stimulated cells were collected, washed, and stained for viability. Fc-block and surface staining for Thy1.1, CD4, and CD8α was done before fixation and permeablization with BD Cytofix/Cytoperm. Intracellular stains (IFNγ and TNFα) were prepared in BD Perm/Wash buffer and incubated with the cells at room temperature for 30 minutes prior to flow cytometry.

### Phosflow staining

T cells were incubated in serum- and cytokine-free conditions for 6 or 12 hours to minimize background phosphorylation events. In the final hour of incubation, 100 ng/mL recombinant human IL-15 (STAT5 phosflow) or 10 ng/mL each of recombinant mouse IFNα and IFNβ (STAT1 phosflow) were added to positive control wells. Cells were collected into 5 mL polypropylene tubes and immediately fixed with warm BD Cytofix, per the manufacturer’s protocol. Samples were permeablized by adding 1 mL of chilled BD Phosflow Perm Buffer III dropwise while vortexing. Phospho-antibody (anti-pSTAT5, anti-pSTAT1) was added directly to samples according to manufacturer’s recommendation and incubated for 1 hr on ice prior to flow cytometry.

### Proliferation assay

Engineered T cells were labeled with CellTrace Violet dye (ThermoFisher Scientific) according to the manufacturer’s protocol. T cells were then co-cultured with CMS5, CMS5r-BCMA, or B16.F10-gp33 tumour cells for 3-5 days at indicated effector:target ratios in T cell media containing 10 ng/mL recombinant human IL-7. Co-cultures were stained with viability dye, Fc-block, and antibodies against CD8α, CD4, and CD90.1 and a defined volume of 123Count eBeads (ThermoFisher Scientific) was added to each sample prior to acquisition by flow cytometry.

Data were analysed using the Proliferation Fit algorithm in FCS Express 7 software. Briefly, data from non-stimulated controls (T cells incubated in media + rhIL-7 alone) were used to set a fixed Starting Generation (i.e. MFI of the “undivided” peak) for each T cell culture. The software then calculated Proliferation Fit Statistics for stimulated samples, including Proliferation Index (the average number of daughter cells that any given T cell in the sample produced). Absolute cell yield was calculated as cell count 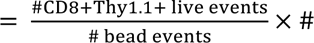 beads loaded into sample.

### In vitro cytotoxicity assay

For luciferase-based cytotoxicity assays, CAR-T cells were co-cultured in triplicate with 5x10^4^ luciferase-expressing target cells at indicated effector:target ratios in a white flat-bottom 96-well plate (Corning) for 12 hours at 37°C. Following co-culture, 0.15 mg/mL D-Luciferin (Perkin Elmer) was added, and luminescence was measured with an open filter using a SpectraMax i3 (Molecular Devices) plate reader. Tumour cell viability was calculated as 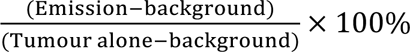.

For imaging-based cytotoxicity assays, P14 CAR-T cells were co-cultured in triplicate with 1x10^4^ NucLight Green-expressing B16.F10-gp33 cells at 4:1, 2:1, and 1:1 E:T ratios for 5 days in an IncuCyte S3 live cell imager (Sartorius). Wells were imaged every 6 hours. The Cell-by-Cell analysis package in the IncuCyte software was used to enumerate live tumour cells in each image based on green fluorescence and cell size. Cell-by-cell data were averaged and normalized to time = 0 for each T cell culture.

### Tumour challenge and combination ACT/OV vaccine therapy

C57BL/6 mice were challenged intradermally with 1x10^5^ B16.F10-gp33 or 2x10^5^ MC38-gp33 cells. One day prior to treatment, tumours were measured, and average tumour burden was normalized across groups. Adoptive transfer occurred at 7-8 days post-tumour inoculation. Cryopreserved T cells were thawed immediately prior to adoptive transfer. Mice received 1x10^6^ CAR+ or an equivalent number of non-engineered T cells in 200 µL of CryoStor® CS10 cryopreservation medium (StemCell Technologies) intravenously. Mice then received 2x10^8^ PFU recombinant VSVΔ51 intravenously at 24 hours post-ACT. For experiments that included systemic IFNAR-1 blockade as part of the therapy, 0.5 mg of anti-mouse IFNAR-1 blocking antibody (clone MAR1-5A3, BioXCell BE0241) in sterile PBS was administered intraperitoneally 16-24 hours prior to vaccination. Tumour volume was calculated as width x length x height. Tumour endpoint was defined as tumour volume ≥1000 mm^3^ or non-resolving tumour ulceration.

### Quantifying peripheral blood T cells after vaccination

At 5 and 12 days following vaccination, a non-terminal blood sample was collected from the submandibular vein as previously described (36). Blood samples were subjected to ACK lysis of erythrocytes prior to peptide stimulation and staining for flow cytometry. The volume of blood collected per animal was calculated as ([final mass of tube + heparin solution + blood sample] - [initial mass of tube + heparin solution])/(blood density = 1.0565 g/mL). A defined volume of 123Count eBeads (ThermoFisher) was added to each PBMC sample after staining, allowing for calculation of absolute T cell counts and a T cell count/volume blood for each animal to be determined following flow cytometric analysis.

### Bioluminescent imaging

Mice that had received either red-shifted luciferase-expressing T cells or red-shifted luciferase-expressing VSVΔ51 as part of the combination therapy protocol described above were anaesthetized with isoflurane and 150mg/kg D-luciferin (Perkin Elmer) was administered intraperitoneally. Animals were monitored for 14 minutes, after which dorsal and ventral images were taken using the IVIS Spectrum imager (Revvity). Total flux signal was quantified with LivingImage v4.2 software (Perkin Elmer).

### RNA and gDNA extraction from murine tissues, PCR and quantitative real-time PCR

Tumours, spleens, and lymph nodes were excised and snap-frozen in liquid nitrogen prior to storage at -80°C. Tissues were homogenized and gDNA and total RNA were collected with a Qiagen AllPrep DNA/RNA/miRNA Universal kit. Reverse transcription of RNA samples was performed using SuperScript IV First-Strand Synthesis System (Invitrogen).

Referencing the original article in which the B16.F10-gp33 cells were generated (29), as well as the article in which the minigene cassette was described (37), we designed a forward PCR primer targeting the region encoding the gp33 peptide and a reverse primer within the published neomycin resistance gene sequence (38). These PCR primers were synthesized by Integrated DNA Technologies (5’ - 3’): AAAGCTGTGTACAATTTCGCCAC and AGAACCTGCGTGCAATCCATC. PCR amplification of the gp33 minigene was performed using a KAPA HiFi PCR Kit (Roche). Primers targeting murine GAPDH were used to generate loading controls for each sample (5’ – 3’): AGGAGCGAGACCCCACTAAC and GGTTCACACCCATCACAAAC. PCR reactions were electrophoresed in a 1% or 3% agarose gels, respectively, and visualized with an Alphaimager (Alpha Innotech).

Quantitative PCR was run using TaqMan Gene Expression Assays (Thermo Fisher Scientific) Hs04982774_m1 (human BCMA/TNFRSF17) and Mm99999915_g1 (murine GAPDH). Data were acquired with a StepOnePlus™ Real-Time PCR System (Applied Biosystems). Relative fold change in expression was determined using the delta/delta CT (2^-ΔΔCT^) method (39), with GAPDH serving as the endogenous control and cDNA from CMSr-BCMA cells as the reference control.

### ELISA

Soluble human BCMA was quantified in cell culture supernatants and serum following infection of CMS5 cells and C57BL/6 mice, respectively, with OV expressing the hBCMA-muFc chimeric antigen. CMS5 cells were infected with indicated multiplicities of infection (MOIs) for 16-24 hours. Supernatants were removed from cell monolayers and passed through a 0.2 µm filter prior to storage at -20°C. Non-terminal submandibular blood samples were collected into Microtainer® SST Blood Collection Tubes (BD), incubated for 30-60 minutes at room temperature, and centrifuged at 3000 xg for 5 minutes at 4°C to isolate serum prior to storage at - 80°C. Undiluted samples (supernatants) and samples diluted 1:100 (serum samples) were analyzed using the R&D DuoSet ELISA human BCMA kit (Cat No. DY193) according to the manufacturer’s protocol. DuoSet ELISA were developed with KPL SureBlue TMB reagent (Seracare) and data were acquired with a SpectraMax i3 (Molecular Devices). Murine IFNα and IFNβ were quantified in serum following ACT/VSV combination therapy in B16.F10-gp33 tumour-bearing C57BL/6 mice. Serum IFNα and IFNβ concentrations were evaluated using Lumikine Xpress mIFNα 2.0 and mIFNβ 2.0 kits (Invivogen) per the manufacturer’s protocol. Serum samples were diluted 1:50 in PBS + 25% FBS and data were acquired with a BioTek Synergy H1 plate reader (Agilent). ELISA data were analyzed with GraphPad Prism (Dotmatics) 4-Parameter Logistic Curve-Fit analysis.

### Statistical analyses

A student’s t-test was used to compare the means of two groups, while one- or two-way analysis of variation (ANOVA) using the Tukey-Kramer multiple comparison test was used to compare three or more groups within an experiment. The Log-rank (Mantel-Cox) test was used to compare survival curves unless otherwise stated. Statistical significance and p values were calculated using GraphPad Prism 10.2.3 for macOS. * p < 0.05, ** p < 0.01, *** p < 0.001, **** p<0.0001, and ns = not significant.

### Study Approval

All animal studies were conducted in compliance with the Canadian Council on Animal Care guidelines and were approved by McMaster University’s Animal Research Ethics Board.

## Results

### Selection and generation of BCMA-specific second-generation CARs for the boosting system

We first sought to generate a murine CAR targeting a known antigen that could subsequently be encoded in recombinant OV vectors. We elected to use known single-chain antibody variable fragments (scFvs) against human antigens to simplify the antigen-CAR pairing. Using well-described scFvs targeting human HER-2, CD19, and BCMA (FRP5, FMC63, and C11D5.3, respectively), we generated four murine CAR constructs with these ScFvs and compared both the surface expression and *in vitro* function of each construct on murine T cells (**Figure 1A**). All four constructs were well-expressed by TCR transgenic T cells and there was distinct separation between CAR+ and CAR-populations (**Figure 1B**). We found that there was an overall decrease in CAR surface expression on day 14 compared to day 6, however this decrease was more substantial in the CD19CAR and CD19WCAR T cells than in the BCMACAR or HER2CAR T cells. Further, a greater proportion of the CD19CAR and CD19WCAR-T cells lost transgene expression altogether by day 14 compared to BCMACAR and HER2CAR-T cells (**Figure 1B**), suggesting that the CD19-targeting CAR constructs may be less stable over time compared with the BCMACAR and HER2CAR constructs. Each construct was able to mediate enhanced cytotoxicity against antigen+ target cells compared to untransduced control cells (**Figure 1C**), demonstrating functionality. We also assayed the cytotoxicity of CAR-expressing DUC18 TCR-transgenic T cells against CMS5 tumour cells, which express the peptide target of the DUC18 TCR (mERK). All cells transduced with the CAR constructs showed TCR-mediated cytotoxicity equivalent to, or better than, untransduced control cells (**Figure 1C**). Based on these data, we chose to go forward with the human BCMA-targeted CAR as our model boosting receptor.

**Figure 1.**
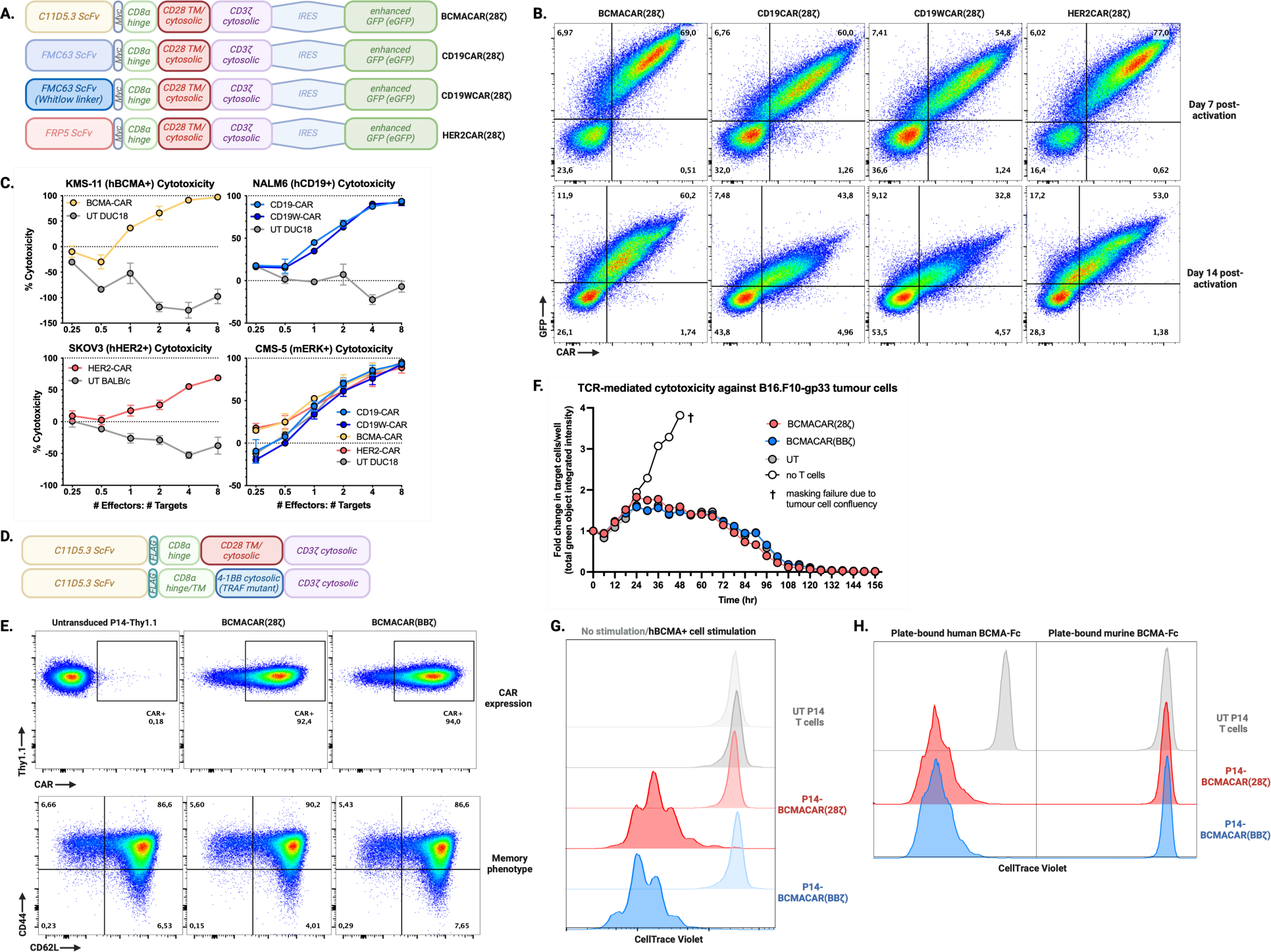
Generation and *in vitro* validation of murine boosting CAR constructs. **A.** Murine CAR-eGFP construct designs. **B.** Flow cytometry analysis of CAR expression in transduced DUC18 T cells at days 6 and 14, detected using target-Fc proteins and anti-human IgG antibody. **C.** Luciferase cytotoxicity assays measuring target cell viability after 12-hour co-culture with CAR-T cells at varying E:T ratios. **D.** Structure of murine CD28ζ and 4-1BBζ second-generation BCMA-targeting CARs. **E.** Day 7 CAR expression and memory phenotype of P14-BCMACAR T cells. See **Figure S1** for gating strategy. **F.** IncuCyte-based cytotoxicity assay of P14 T cells against NucLight Green B16.F10-gp33 cells, normalized to baseline. **G.** CTV dilution assay showing proliferation of BCMACAR-P14 or UT P14 T cells after 4-day co-culture with CMS5r-BCMA cells. **H.** CTV dilution of P14 BCMACAR-T cells after 72-hour culture with plate-bound human or murine BCMA-Fc.

We next engineered the BCMA-specific scFv onto a 4-1BB-based second-generation CAR backbone (BCMACAR-BBζ) in addition to the CD28-based CAR backbone (BCMA-28ζ) (**Figure 1D**). As we have previously established that a less-differentiated central (T_cm_) or stem cell (T_scm_) memory phenotype is critical in the observed synergy between adoptively transferred T cells and VSV booster vaccines (16), we also confirmed that these BCMACAR T cells demonstrate a predominantly (>90%) T_cm_/T_scm_ phenotype at the end of *ex vivo* culture (**Figure 1E; gating is shown Figure S1**). P14 TCR-transgenic T cells engineered with the BCMACAR constructs also showed TCR-mediated cytotoxicity against tumour cells bearing cognate gp33 antigen comparable to non-engineered P14 cells (**Figure 1F**) and robust proliferation upon CAR engagement (**Figure 1G**)

Finally, we confirmed that the C11D5.3 ScFv in our BCMACARs shows no cross-reactivity with murine BCMA (**Figure 1H**), as this could lead to unwanted engagement of endogenous mBCMA-expressing cells *in vivo* potentially leading to toxicities and vaccine-independent T cell expansion.

### Boosting of the CAR-engineered T cells with a paired rhabdovirus vaccine

For the paired boosting antigen, we elected to evaluate two forms of signaling-incompetent human BCMA: (i) surface-expressed BCMA fused to a short truncated human NGFR cytosolic domain (hBCMAtNGFR) and (ii) secreted as an Ig-fusion protein (hBCMA-Fc) (**Figure S2 and Figure S3**). To evaluate the system in tumour-bearing animals, we first treated B16.F10-gp33 tumour-bearing C57BL/6 mice with P14 BCMACAR(28ζ), BCMACAR(BBζ), or untransduced (UT) T cells in combination with VSV-hBCMAtNGFR; the frequency of transferred cells in the blood was evaluated at early and late timepoints and tumour growth was monitored (**Figure 2A; gating is shown in Figure S4**). We observed expansion of the transferred T cells at day 5 post-vaccination only in those animals that received BCMACAR-T cells in combination with the paired VSV-hBCMAtNGFR vaccine (**Figure 2B**). The frequency of transferred cells observed in the blood at the early timepoint was variable across individual mice and the CD28-based CAR construct drove more robust expansion compared to the 4-1BB CAR. We also observed CAR downregulation on the surface of the boosted cells, consistent with robust T cell activation (**Figure 2C**). However, the boosted BCMACAR-T cells showed poor persistence in vaccinated animals, regardless of which co-stimulatory domain was used (**Figure 2B**). Despite the transient persistence of the boosted cells, the combination of the boosting CD28-CAR and boosting vaccine led to short-term tumour regressions and an overall survival benefit compared to treatment with either CAR-T cells alone or non-engineered T cells with the booster vaccine (**Figure 2D, Figure S5**).

**Figure 2.**
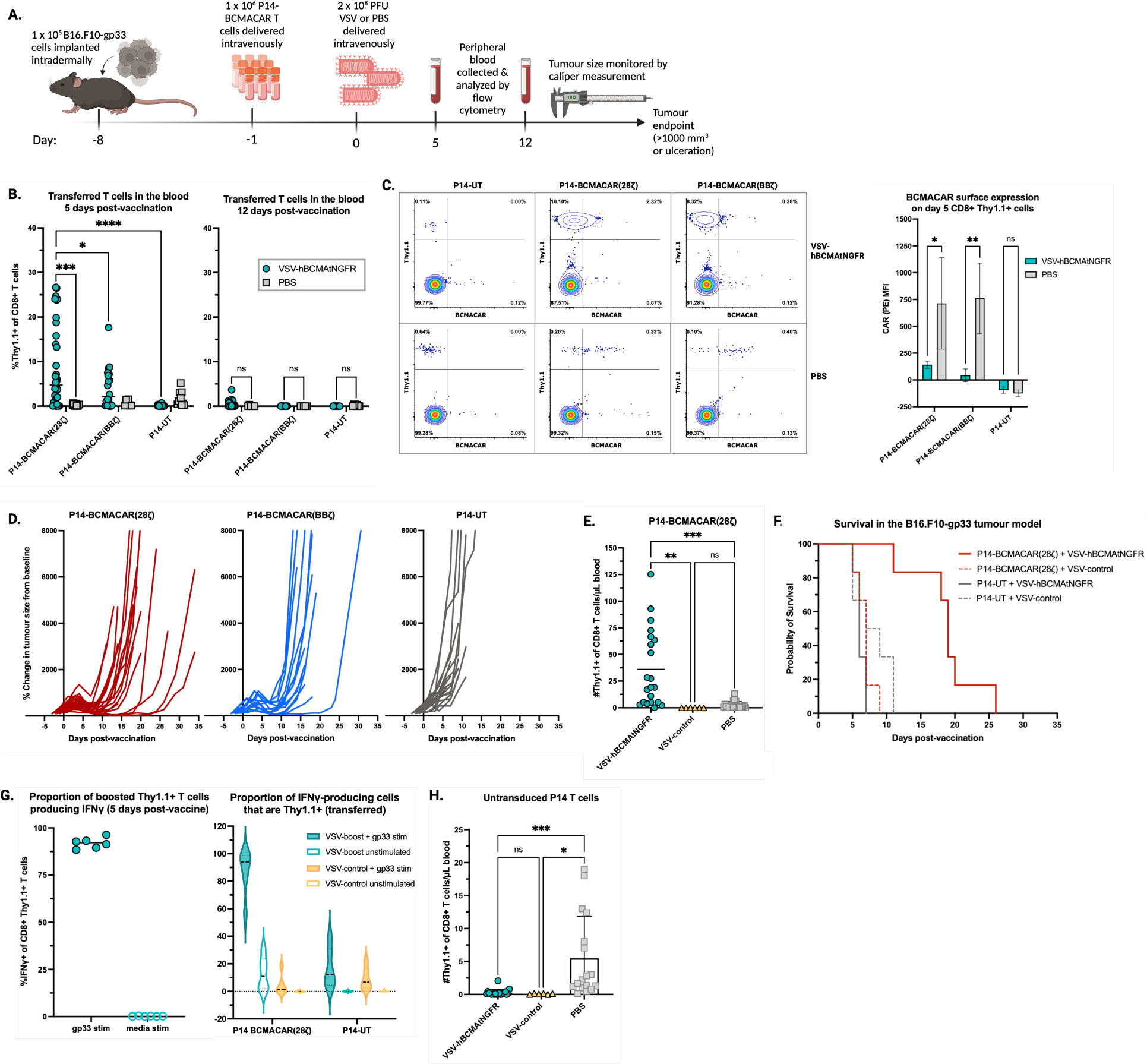
CAR-T cell expansion is robust, but transient, following VSV vaccination. **A.** 1x10^6^ P14 CAR-T/T cells were given i.v. 8 days post-tumour implantation, followed by 2x10^8^ PFU VSV-hBCMAtNGFR/control 24 hours later. **B-C.** Flow cytometric analysis of blood at days 5 and 12 post-vaccination, evaluating viability and surface markers (CD8, CD4, Thy1.1, BCMACAR). See **Figure S4** for gating strategy. **D.** Tumour volume measured by caliper every 2-3 days, normalized to pre-treatment size. Truncated curves indicate tumour ulceration. Data in B-D are representative of 9 independent experiments. See **Figure S5** for the corresponding survival curves. **E, G-H.** PBMCs were analyzed for antigen reactivity via IFNγ production after gp33 stimulation, with absolute counts determined using counting beads. **F.** Survival post-vaccination with either VSV-hBCMAtNGFR or VSV-control.

We next assessed whether the VSV vaccine encoding the BCMA-Fc fusion (VSV-hBCMAmFc) could provide comparable, or enhanced, boosting relative to VSV-hBCMAtNGFR. After confirming that the hBCMAmFc fusion protein was expressed, albeit transiently, *in vivo* following vaccination (**Figure S6 A, B**), we evaluated the ability of VSV-hBCMAmFc to boost P14-BCMACAR T cells *in vivo* (**Figure S6C**). Although robust boosting of the transferred cells was achieved following TCR engagement with VSV-gp33, we did not observe any expansion of the P14-BCMACAR T cells with the soluble antigen vaccine (**Figure S6D**), suggesting that the soluble antigen may not be sufficient to engage the transferred cells. As such, all further work employed VSV-hBCMAtNGFR as the boosting vector.

To confirm the requirement of VSV-expressed target antigen, P14 BCMA-CAR28ζ cells were transferred into wild type recipients followed by immunization with VSV-hBCMAtNGFR or VSV expressing an irrelevant antigen (VSV-control). P14-BCMACAR T cell expansion only occurred in the context of paired VSV-hBCMAtNGFR vaccination (**Figure 2E**). Moreover, when compared to the frequency of non-engineered P14 T cells (P14-UT) in the blood of non-vaccinated (PBS control) mice, there were significantly fewer transferred cells 5 days post-vaccination when mice were treated with VSV, further demonstrating that in the absence of CAR or TCR engagement transferred T cells fail to engraft and expand in the presence of VSV (**Figure 2E**). No survival benefit was observed in the absence of T cell expansion, demonstrating that efficacy is dependent on synergy between the P14 CAR-T cells and the paired VSV booster vaccine rather than either the T cells or virus independently (**Figure 2F**). We also evaluated the ability of the boosted P14 T cells to respond to stimulation through their endogenous TCR at day 5 post-vaccination by pulsing the PBMCs with gp33 peptide and evaluating cytokine production. Nearly all boosted P14 BCMACAR-T cells produced IFNγ in response to *ex vivo* stimulation with gp33 peptide, indicating that TCR-mediated anti-tumour function remains intact following CAR-driven *in vivo* expansion (**Figure 2G**). Importantly, we observed that mice that had received VSV-control showed complete attrition of the transferred BCMACAR-T cells by day 5 post-vaccination (**Figure 2H)**, suggesting that the presence of VSV may be inhibiting T cell engraftment and expansion altogether in the absence of T cell engagement.

### Transient IFNAR1 blockade during VSV vaccination enhances CAR-T cell expansion and extends tumour regression

Type I interferons (IFN-I) are known to cause attrition of memory T cell in the early stages of virus infection (40). Considering the data in Figure 2E, we hypothesized that VSV-induced IFN-I was preventing efficient engraftment and expansion of transferred T cells. To address this possibility, we administered a single dose of anti-IFNAR1 blocking antibody between ACT and VSV vaccination in B16.F10-gp33 tumour-bearing animals (**Figure 3A**). Compared to mice that received either PBS or a matched isotype control antibody, the mice that received IFNAR1 blockade showed a robust enhancement in P14-BCMACAR T cell boosting at day 5 post-vaccination (**Figure 3B**). Of note, the combination of VSV vaccine and IFNAR blockade not only led to enhanced expansion of the adoptively transferred cells, but also prevented VSV-driven attrition of endogenous CD8+ T cells in mice that received untransduced T cells (**Figure 3C**). The T cells boosted in the presence of IFNAR blockade also showed reduced CAR downregulation at this early time point (**Figure 3D**). Despite the increase in expansion of transferred CAR-T cells at day 5 post-vaccination, IFNAR1 blockade did not enhance persistence of these cells (**Figure 3E**). IFNAR1 blockade in combination with CAR-T cells and VSV vaccination enhanced survival of tumour-bearing mice compared to those that received the isotype control (**Figure 3F**); however, all tumours relapsed with continued expression of gp33 antigen (**Figure 3G**). These data confirmed that P14 CAR-T cell boosting is hindered by type I IFN signaling following VSV vaccination, but the boosted cells are unable to mediate complete clearance of antigen-positive cells.

**Figure 3.**
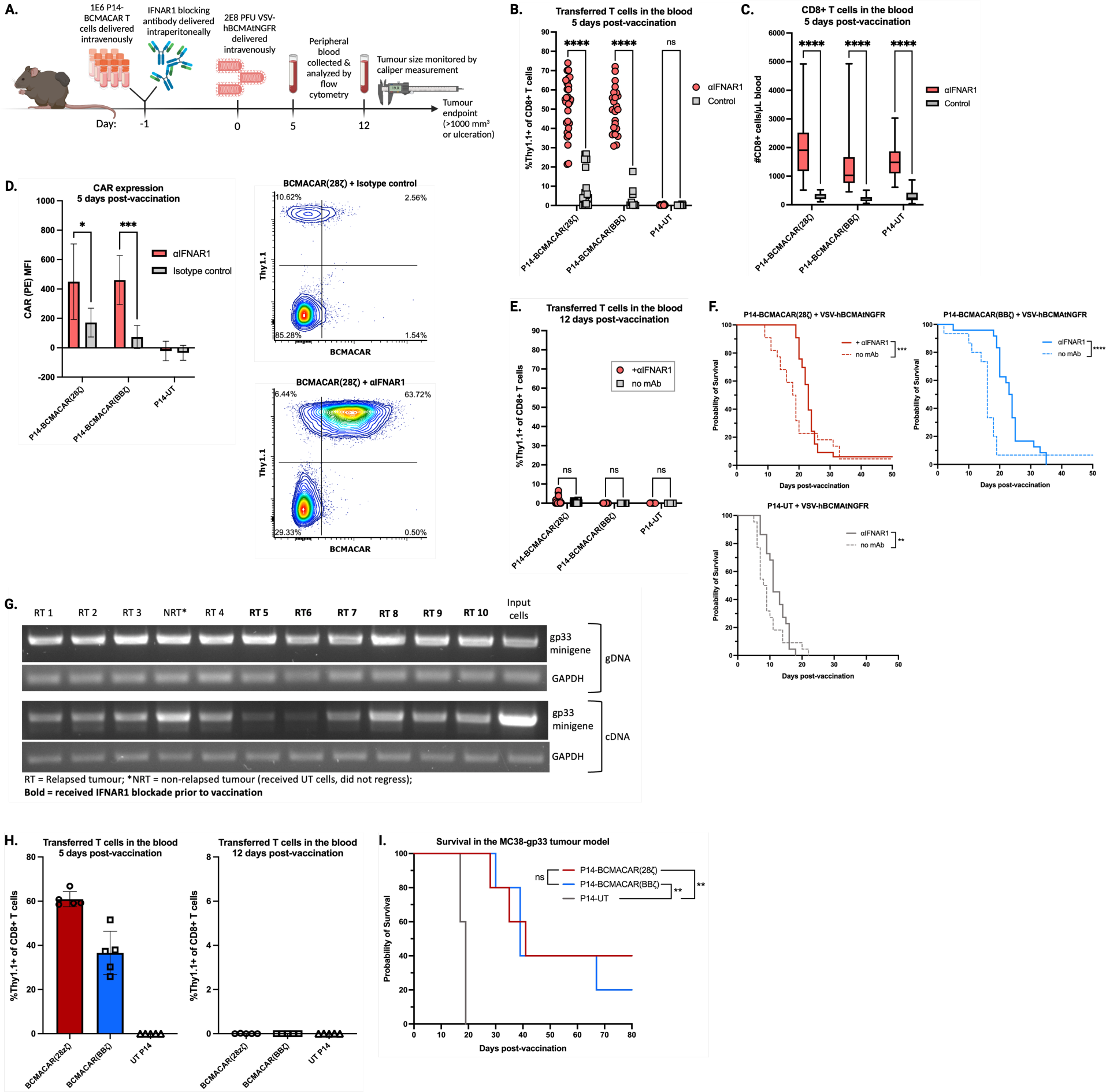
Transiently blocking type I IFN signaling during vaccination enhances transferred CAR-T cell expansion and efficacy. **A.** 1x10^6^ P14 CAR-T/T cells were given i.v. 8 days post-tumour implantation, followed by 2x10^8^ PFU VSV-hBCMAtNGFR/control 24 hours later. Mice were given αIFNAR1 blocking antibody or isotype control antibody intraperitoneally 16-22 hours prior to vaccination. **B-E.** Flow cytometric analysis of blood at days 5 and 12 post-vaccination, evaluating viability and surface markers (CD8, CD4, Thy1.1, BCMACAR). **F.** Survival analysis of B16.F10-gp33 tumour-bearing mice (representative of 8 independent experiments, Gehan-Breslow-Wilcoxon test). **G.** The gp33 minigene was PCR-amplified from genomic DNA and cDNA of relapsed B16.F10-gp33 tumours following adoptive transfer and VSV vaccination ± αIFNAR1. GAPDH was amplified in the same samples as a housekeeping control. Genomic DNA and cDNA from minimally cultured B16.F10-gp33 cells were used as input control samples. **H.** Expansion and persistence of boosted P14 T cells in the presence of IFNAR1 blockade in MC38-gp33 tumour-bearing recipients. **I.** Survival of MC38-gp33 tumour-bearing mice following vaccination in the presence of IFNAR1 blockade (representative of 2 independent experiments, log-rank Mantel-Cox test).

Recognizing that the IFN-I effect may be influencing the performance of VSV-BCMAmFc boost vaccine, we re-evaluated this vector in the presence of IFNAR1 blockade and assessed BCMACAR-T cell boosting and anti-tumour efficacy (**Figure S7A**). However, again, there was no expansion of transferred BCMACAR-T cells upon VSV-hBCMAmFc boosting, even in the presence of IFNAR1 blockade (**Figure S7B**). As a result, this treatment combination led to no anti-tumour efficacy and no improvement to survival in the B16.F10-gp33 model **(Figure S7C, D**). The marked difference in outcome between the membrane bound BCMA and the BCMA-Fc fusion suggest that stable cell membrane association of the boosting antigen is required for effective CAR-mediated T cell boosting in this setting.

To evaluate whether these findings extend to a different syngeneic tumour model, we assessed the same treatment strategy in the MC38-gp33 murine colon carcinoma model (**Figure 3H and 3I**). In the presence of IFN-I blockade, P14-BCMACAR T cells underwent robust expansion upon VSV-hBCMAtNGFR vaccination in MC38-gp33 tumour-bearing mice, comparable to that observed in the B16.F10-gp33 model (**Figure 3H**). Also similarly, the transferred P14 T cells did not persist past initial expansion, demonstrating that the lack of persistence following boosting is not a model-specific phenomenon (**Figure 3H**). There was also a survival benefit to the combination treatment in the MC38-gp33 model (**Figure 3I**) confirming the utility of this expansion strategy in a secondary model.

### IFN-I suppresses boosting in a T cell-extrinsic manner

We hypothesized that IFN-I may be directly suppressing CAR-mediated T cell proliferation. To address this, we performed *in vitro* proliferation assays with P14-BCMACAR T cells stimulated with gp33-expressing or BCMA-expressing tumour cells in the presence, or absence, of IFNα or IFNβ (**Figure 4A and 4B**). Neither IFNα nor IFNβ (10 or 25 ng/mL) suppressed the proliferation of P14 T cells following stimulation through the CAR (**Figure 4B**). In contrast, a dose-dependent increase in proliferation was observed following TCR stimulation in the presence of either IFNα or IFNβ, likely driven by IFN-mediated MHC upregulation on the B16.F10-gp33 cells.

**Figure 4.**
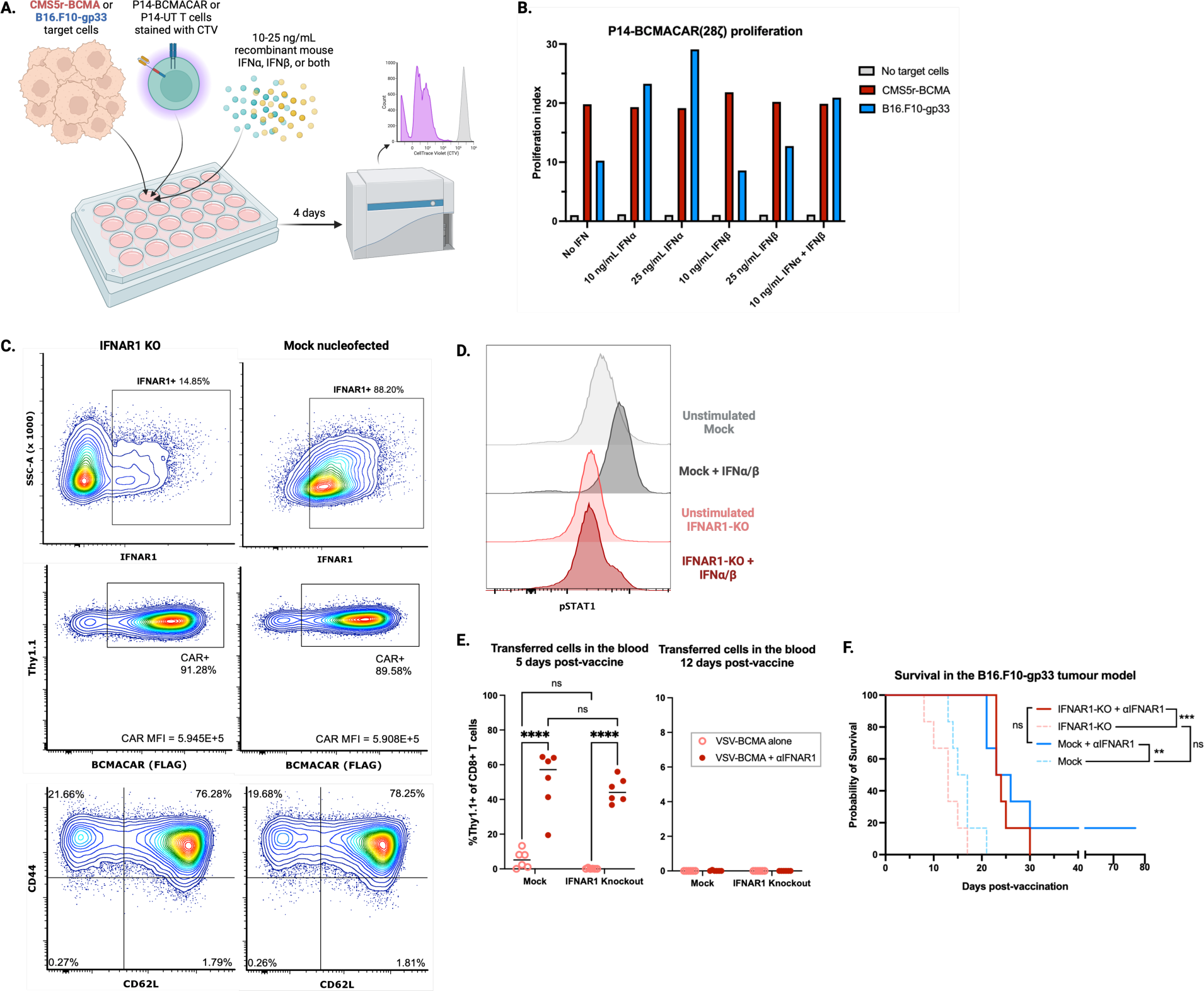
Type I interferon-mediated suppression of CAR-T cell boosting is mediated through a T cell-extrinsic mechanism. **A.** CellTrace Violet dilution analysis of P14-BCMACAR T cell proliferation after CAR(CMS5r-BCMA)/TCR (B16.F10-gp33) stimulation ± mIFNα/β. Proliferation data are representative of two biological replicates. **B.** Proliferation index of live CD8+ Thy1.1+ cells following coculture. **C.** Flow cytometry analysis of IFNAR1-KO and mock-nucleofected P14-BCMACAR(28ζ) cells at day 5 post-activation, evaluating IFNAR1, CD8, Thy1.1, CD62L, CD44, and CAR (FLAG) expression. MFI denotes mean (arithmetic) fluorescence intensity of the FLAG marker in the CAR+ population. **D.** pSTAT1 quantification after 1h IFNα/β stimulation in control vs IFNAR1-KO P14-BCMACAR(28ζ) T cells. **E.** Analysis of transferred Thy1.1+ CD8+ T cells in blood at days 5 and 12 post-vaccination in B16.F10-gp33 tumour model: 1x10^6^ IFNAR1-KO/mock CAR-T cells were given i.v., ± αIFNAR1 at 2-4h, followed by VSV-BCMA at 24h. **F.** Survival of B16.F10-gp33 tumour-bearing mice post-VSV-BCMA vaccination (representative of 2 independent experiments, log-rank Mantel-Cox test).

To address the possible effects of IFN-I on CAR-T cells *in vivo*, we edited out endogenous murine IFNAR1 using CRISPR/Cas9 and achieved approximately 80% knock-down without impacting transduction, CAR expression, or memory phenotype (**Figure 4C**). The IFNAR1-KO P14-BCMACAR(28ζ) T cells have lower baseline STAT1 phosphorylation compared to mock-nucleofected control cells and fail to induce STAT1 phosphorylation following stimulation with IFNα and IFNβ (**Figure 4D**). These results confirmed that our edited CAR-T cells were phenotypically and functionally defective in IFNAR1 expression and signaling.

Mock-nucleofected (Mock) control and IFNAR1-KO P14-BCMACAR(28ζ) T cells were adoptively transferred into B16.F10-gp33 tumour-bearing mice and boosted with VSV-hBCMAtNGFR in the presence or absence of αIFNAR1 blocking antibody (**Figure 4E and 4F**). Here, we observed robust CAR-T cell boosting only in the mice that received IFNAR1 blockade (**Figure 4E**). Interestingly, in the absence of systemic IFNAR1 blockade, the Mock cells showed greater engraftment and expansion at the early timepoint compared to the IFNAR1-KO cells. Regardless of expansion on day 5 post-vaccination, there was no persistence of boosted cells by day 12 post-vaccination (**Figure 4E**). The Mock and IFNAR1-KO T cells performed similarly in terms of anti-tumour efficacy, with the mice that received IFNAR1 blockade showing prolonged tumour control and survival compared to those that received no blocking antibody (**Figure 4F**). These data confirmed that type I interferon-mediated suppression of CAR-T cell expansion following VSVΔM51 vaccination is mediated through a T cell-extrinsic mechanism.

### Blocking type I IFN signaling during vaccination increases duration and magnitude of boosting antigen load

A potential T cell-extrinsic mechanism by which type I IFN could be influencing CAR-T cell expansion in our system is through premature clearance of the VSV vaccine, thus limiting the availability of antigen to stimulate the CAR. We observed that peak expression of the hBCMAtNGFR antigen by VSVΔM51 occurred in the spleen, lymph nodes, and tumour around 6 hours post-vaccination in the absence of BCMACAR-T cells and IFNAR1 blockade (**Figure 5A**). To determine whether IFNAR1 blockade may be changing antigen expression patterns, we first used serum levels of the hBCMAmFc antigen (**Figure S3, Figure S6**) as a measure of VSV expression in the presence/absence of IFNAR1 blockade. Indeed, serum hBCMAmFc levels displayed an exponential increase in the presence of IFNAR1 blockade and a shift in peak serum levels from 6 hours to 48 hours post-infusion (**Figure S8**). To directly assess whether IFNAR1 blockade similarly extends the expression of hBCMAtNGFR, we employed a VSV vaccine (VSV-rsLuc/hBCMAtNGFR) that encodes both the hBCMAtNGFR antigen and red-shifted luciferase (rsLuc) transgenes linked as a single polyprotein through a 2A sequence such that luciferase serves as a direct measure of antigen expression (**Figure S9A**). We confirmed that VSV-rsLuc/hBCMAtNGFR demonstrates transgene expression levels and kinetics comparable to the original single-expression vaccine vector, and that the luminescent signal from this virus can be detected *in vivo* post-vaccination (**Figure S9B-D**). B16.F10-gp33 tumour-bearing mice were given P14-BCMACAR(28ζ) T cells followed by VSV-rsLuc/hBCMAtNGFR in the presence or absence of IFNAR1 blockade (**Figure 5B**).We observed that VSV transgene expression was increased at all timepoints when vaccination was given in the presence of IFNAR1 blockade (**Figure 5B**). P14 T cell expansion and anti-tumour efficacy in this experiment were consistent with what we have observed with the single-expression VSV-hBCMAtNGFR vaccine in the presence of IFNAR1 blockade (**Figure 5C, D**). Finally, we evaluated serum levels of IFNα and IFNβ in tumour-bearing mice following VSV-hBCMAtNGFR boosting of P14-BCMACAR(28ζ) T cells in the presence and absence of IFNAR1 blockade. Similar to the antigen expression patterns we observed, IFNα and IFNβ concentrations peaked at 6 hours post-vaccination, declined at 24 hours, and were below detectable levels from 48 hours onwards (**Figure 5E**). Therefore, transient IFNAR1 blockade overcomes the initial burst of type I IFN production immediately following VSV vaccination to allow for robust boosting antigen expression at these early timepoints. Taken together, these data indicate that IFNAR1 blockade in our system is acting to increase boosting antigen availability in the days following vaccination.

**Figure 5.**
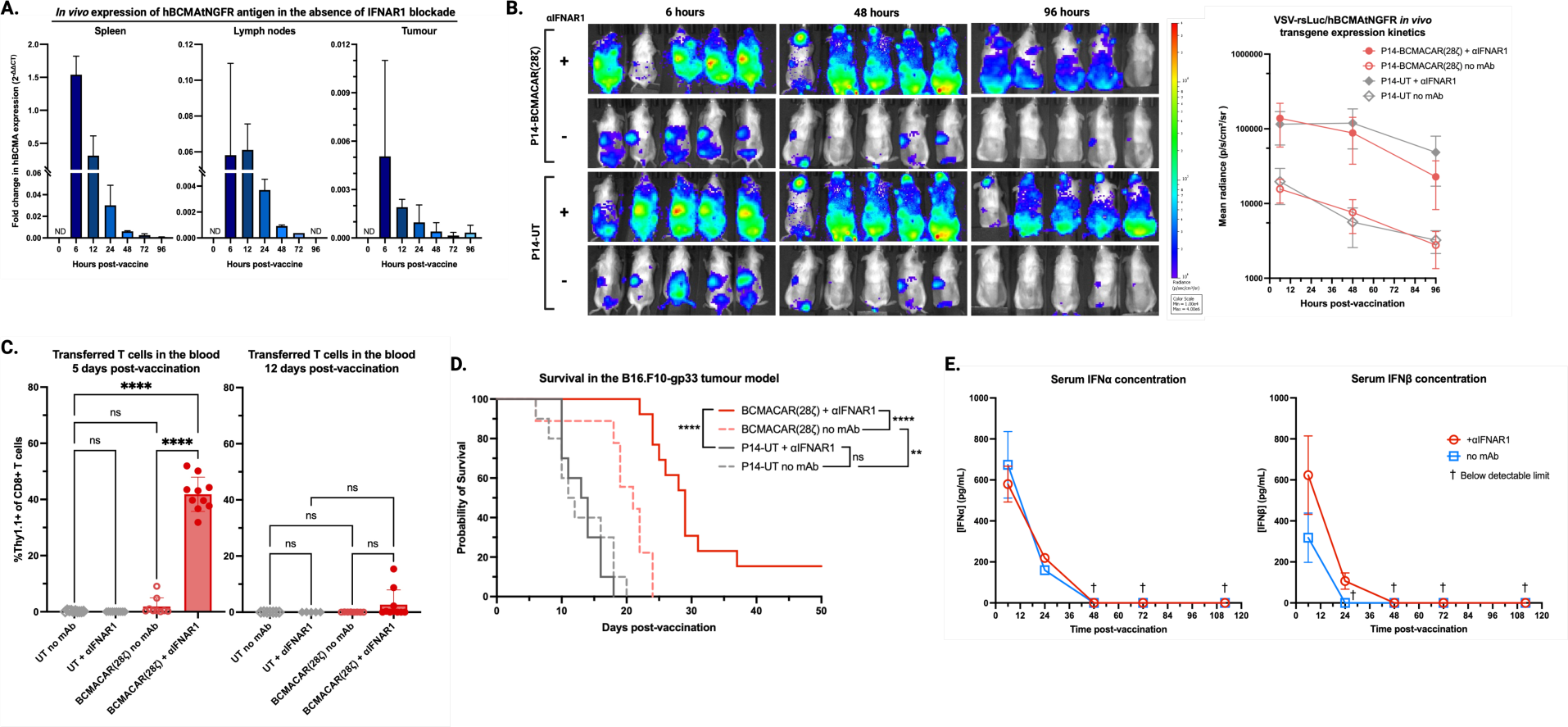
Systemic IFNAR1 blockade enhances boosting antigen expression in vaccinated animals. **A.** qPCR analysis of human BCMA expression in tissues isolated from B16.F10-gp33 tumour-bearing mice at various times post-VSV-hBCMAtNGFR vaccination, normalized to GAPDH with CMS5r-BCMA cDNA as reference control. **B.** IVIS analysis of VSV transgene expression over time following VSV-rsLuc/hBCMAtNGFR vaccination in B16.F10-gp33-tumour bearing C57BL/6 nude mice treated with P14-BCMACAR/UT T cells ± αIFNAR1. **C.** Analysis of transferred Thy1.1+ CD8+ T cells in the blood at days 5 and 12 post-vaccination. See **Figure S9** for VSV-rsLuc/hBCMAtNGFR vaccine validation. **D.** Survival of B16.F10-gp33 tumour-bearing mice post-VSV-BCMA vaccination (representative of 2 independent experiments, log-rank Mantel-Cox test). **E.** Serum IFNα/β concentrations at various timepoints after treatment with P14 BCMACAR-T cells, αIFNAR1, and VSV-hBCMAtNGFR (n=4 biological replicates, run in technical triplicate).

### Boosting through the TCR with tumour associated antigen leads to improved persistence of transferred T cells and complete eradication of antigen-expressing tumour cells

Our prior work demonstrated that boosting P14 T cells through their TCR using VSV-gp33 could promote tumour clearance without requiring IFNAR blockade (17). Those studies employed a different T cell manufacturing method, which may influence T cell performance. Therefore, to assess whether boosting through the TCR might change the behaviour of the BCMACAR(28ζ)-engineered P14 T cells, we infused mice with engineered T cells, administered anti-IFNAR1 and boosted with either VSV-hBCMAtNGFR, to boost via the CAR, or VSV-gp33, to boost through the TCR. The data revealed that both strategies resulted in similar expansion at day 5 post-vaccine but only boosting through the TCR resulted in marked persistence at day 12 (**Figure 6A**). Both boosting strategies resulted in a comparable therapeutic effect where the tumours initially regressed but subsequently relapsed (**Figure 6B**). The improved therapeutic efficacy could be due, in part, to the engagement of endogenous T cells, as naturally occurring gp33-reactive T cells also expanded in the mice receiving VSV-gp33 (**Figure 6C**). A key difference is revealed through the analysis of the target antigen in the relapsed tumours. Whereas the relapsed tumours expression in mice boosted with VSV-hBCMAtNGFR retained gp33 antigen, the relapsed tumours in mice boosted with VSV-gp33 were devoid of the gp33 antigen gene both in the presence and absence of IFNAR1 blockade (**Figure 6D**).

**Figure 6.**
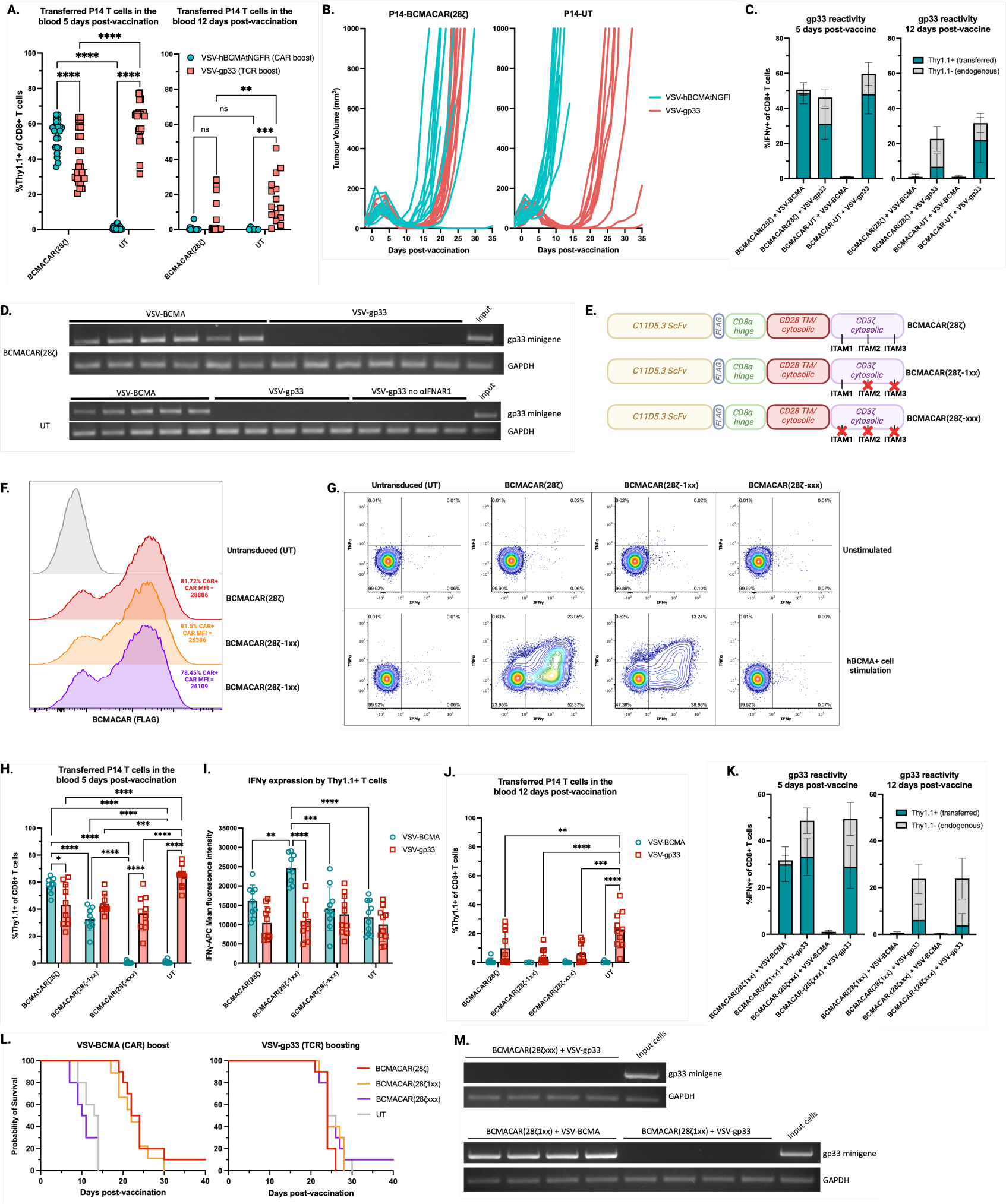
CAR signaling capacity does not impact persistence or antitumour efficacy of the engineered T cells following VSV boosting. 1x10^6^ P14 BCMACAR-T/UT cells were administered to B16.F10-gp33 tumour-bearing C57BL/6 recipients, followed by αIFNAR1 and 2x10^8^ PFU VSV-hBCMAtNGFR or VSV-gp33. **A, C.** 5 and 12 days after vaccination, non-terminal bleeds were collected from the facial vein of each mouse. Following ACK lysis to remove erythrocytes, cells were incubated in the presence of gp33 peptide for 1 hour alone followed by 4 hours in the presence of Brefeldin A to stimulate antigen-reactive CD8+ T cells. Prior to flow cytometric analysis, cells were stained with a fixable viability dye, a surface antigen cocktail (αCD8, αCD4, αThy1.1, and αFLAG [CAR]), and an intracellular stain against IFNγ. **B.** Tumour volume (L×W×H) was measured every 2-3 days; truncated curves indicate tumour ulceration prior to 1000mm^3^. Representative of 3 independent experiments. **D, M.** PCR analysis of gp33 minigene and GAPDH in gDNA from relapsed tumours. **E.** Design of BCMACAR(28ζ1xx) and BCMACAR(28ζxxx) variants with Y>F mutations in the two distal or all three CD3ζ ITAMs. **F.** CAR expression and **G.** cytokine production after CAR (CMS5r-BCMA) stimulation in P14 T cells. **H-K.** *In vivo* CAR-T cell expansion and gp33 antigen reactivity following VSV-hBCMAtNGFR or VSV-gp33 vaccination were evaluated as in (**A.**). Data for BCMACAR(28ζ) and UT P14 T cells were reproduced from (**Α.**) to facilitate visual comparison. **L.** Survival of B16.F10-gp33 tumour-bearing mice following VSV vaccination (representative of 2 independent experiments). See **Figure S11** for the corresponding tumour growth curves.

### Tuning CAR surface expression and signal strength does not lead to improved T cell functionality following boosting

CAR surface expression is positively correlated with tonic (antigen-independent) signaling, meaning that high levels of receptor expression often lead to exhaustion and dysfunction of CAR-T cell products (41). Indeed, a recent report correlated CAR-T cell dysfunction with IFN-dependent upregulation of CAR expression (42). We hypothesized that high CAR expression could be driving poor CAR-T cell persistence. We found that dual-expression CAR constructs containing an IRES and Thy1.1 transgene downstream of the CAR (**Figure S10A**) led to greatly reduced CAR expression on T cells compared to single-expression CAR constructs, despite similar transduction efficiencies, memory phenotype, and *in vitro* proliferation upon CAR or TCR stimulation (**Figure S10B, C**). Boosting of CAR(lo) T cells was associated with improved persistence in fraction of the treated animals relative to the CAR(hi) T cells, but this did not achieve significance (**Figure S10D**). Moreover, there was no different in therapeutic efficacy between the CAR(lo) and CAR(hi) T cells (**Figure S10E**).

Given that difference in receptor levels can confound the interpretation of these experiments, we elected to attenuate CAR signaling by removing 2 ITAMs from the CD3ζ domain, as described previously (43), to yield BCMACAR(28ζ1xx). T cells engineered with BCMACAR(28ζ1xx) displayed CAR expression comparable to BCMACAR(28ζ). As an additional control for the influence of CAR signaling on T cell performance, we generated a CAR devoid of ITAMs, named BCMACAR(28ζxxx) which also displayed surface expression comparable to BCMACAR(28ζ).

T cells engineered to express BCMACAR(28ζxxx) demonstrated no boosting when exposed to VSV-hBCMAtNGFR, confirming the role of the CD3ζ domain in the antigen-mediated boost (**Figure 6H**). Boosting of engineered T cells through BCMACAR(28ζ1xx) resulted in reduced primary expansion at day 5 post-vaccine relative to T cells boosted through BCMACAR(28ζ), consistent with attenuated signaling (**Figure 6H**). The T cells boosted through BCMACAR(28ζ1xx) also displayed higher functionality at day 5 relative to T cells boosted through BCMACAR(28ζ), as defined by the per-cell level of IFN-gamma production (**Figure 6I**). Despite these effects, the T cells boosted through BCMACAR(28ζ1xx) demonstrated no marked benefit to persistence (**Figure 6J**) and therapeutic efficacy was comparable to T cells boosted through BCMACAR(28ζ) (**Figure 6L, Figure S11)**.

In contrast, persistence was only observed when the T cells were boosted through the TCR, and non-engineered T cells showed persistence more consistently and to a greater level than the engineered cells, regardless of which CAR was expressed (**Figure 6J**). Thus, poor persistence may be both intrinsic to the engineered T cells as well as a feature of the mechanism of boosting (i.e. CAR vs. TCR). As observed earlier, only VSV-gp33 vaccination was associated with expansion of endogenous gp33-reactive T cells and loss of the gp33 antigen in relapsed tumours, irrespective of CAR signaling capacity (**Figure 6K, M**).

### Introduction of an IL-15 transgene fails to improve persistence of T cells following CAR-mediated boosting

Several groups have shown that IL-15 coexpression can improve CAR-T cell engraftment, persistence, and anti-tumour efficacy in pre-clinical solid tumour models (44–47). This considered, we inserted the native murine IL-15 sequence downstream of our BCMACAR(28ζ) construct following an IRES (**Figure 7A**). Indeed, T cell transduction efficiency and CAR expression of the single-expression BCMACAR(28ζ) and the dual-expression BCMACAR(28ζ)/mIL-15 construct were comparable (**Figure 7B**). We also observed an increase in frequency of T_eff_/T_em_ populations in cultures engineered with BCMACAR(28ζ)/mIL-15 when compared to BCMACAR(28ζ) alone, consistent with increased STAT5 signaling (**Figure 7B**). Baseline pSTAT5 levels of BCMACAR(28ζ) T cells were comparable to non-engineered (UT) T cells, whereas BCMACAR(28ζ)/mIL-15 cells showed increased pSTAT5 relative to both (**Figure 7C**). When evaluated in an *in vitro* proliferation assay, proliferation index was comparable between the single- and dual-expression cells following CAR and TCR stimulation, but the BCMACAR(28ζ)/mIL-15 cells showed increased viability following CAR stimulation at the end of the co-culture (**Figure 7D**). These data suggest that the IL-15 transgene is functional and can promote T cell survival following CAR-mediated engagement and expansion *in vitro*.

**Figure 7.**
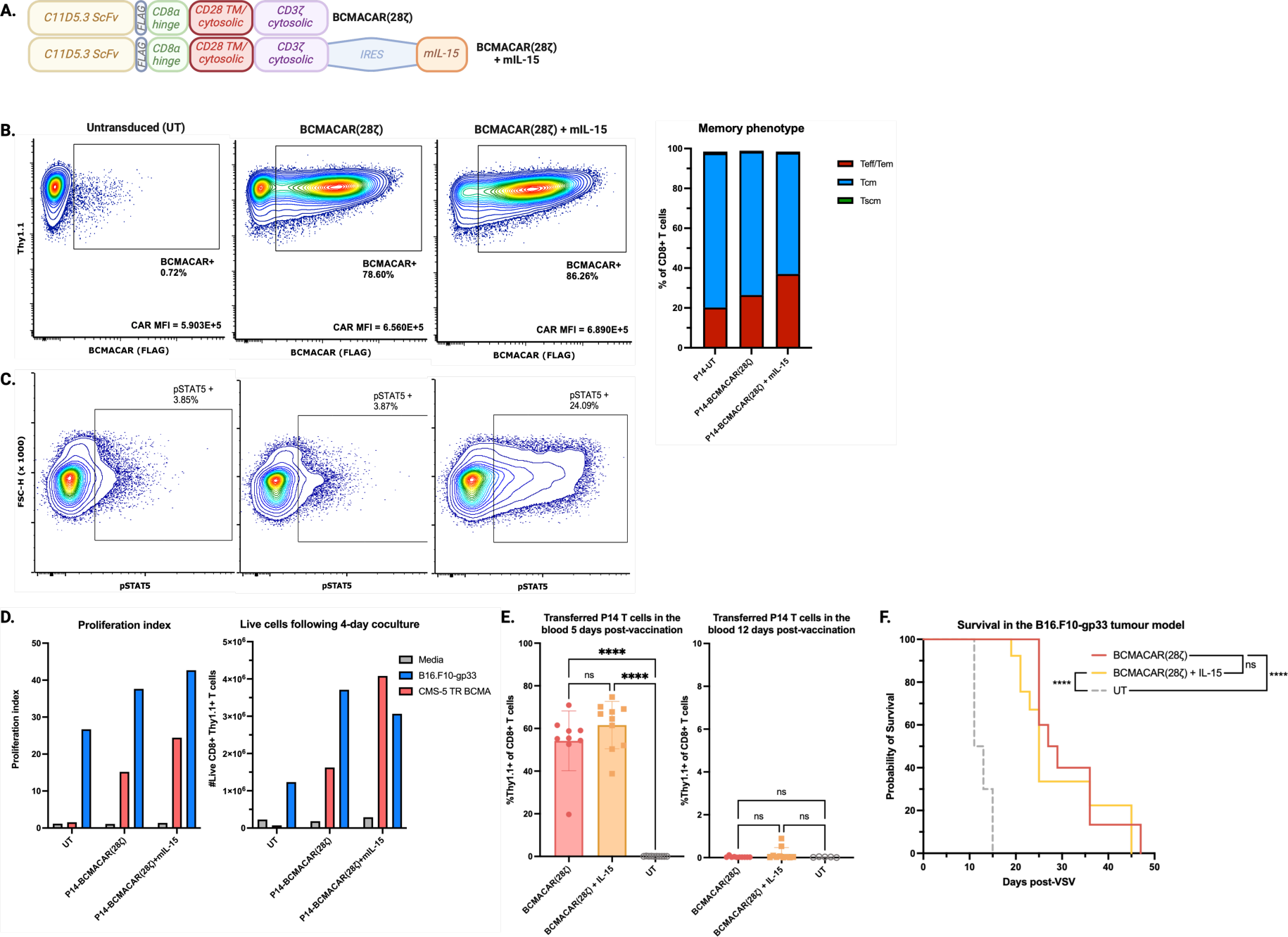
Co-expression of an IL-15 transgene in BCMACAR-T cells is not sufficient to enhance *in vivo* persistence and antitumour efficacy following VSV boosting. **A.** Design of single BCMACAR(28ζ) and dual BCMACAR(28ζ)/mIL-15 constructs. **B.** Phenotypic analysis of day 7 P14-BCMACAR/mIL-15 T cells for CD8, Thy1.1, CD62L, CD44, and FLAG (CAR) expression. MFI denotes mean (arithmetic) fluorescence intensity of the FLAG marker in the CAR+ population. **C.** pSTAT5 (pY694) analysis by flow cytometry after 12h serum/cytokine starvation. **D.** Proliferation index and absolute counts of P14 T cells after 4-day co-culture with CMS5r-BCMA or B16.F10-gp33 targets, measured by CTV dilution. **E.** B16.F10-gp33 tumour-bearing C57BL/6 mice were treated with P14-BCMACAR T cells, αIFNAR1 (4h post-ACT), and VSV-hBCMAtNGFR (24h post-ACT). Blood was collected on days 5 and 12 post-vaccination and evaluated for the presence of CD8+ Thy1.1+ transferred P14 T cells. **D.** Survival of B16.F10-gp33 tumour-bearing animals following vaccination (representative of 2 independent experiments, log-rank Mantel-Cox test). See **Figure S12** for the corresponding tumour growth curves.

We next transferred the single- and dual-expression P14 CAR-T cells into B16.F10-gp33 tumour-bearing mice and evaluated expansion and persistence following VSV-hBCMAtNGFR boosting (**Figure 7E, F**). Both the BCMACAR(28ζ) and BCMACAR(28ζ)/mIL-15 T cells showed robust and comparable expansion in peripheral blood at day 5 post-vaccination, demonstrating that the IL-15 transgene does not inhibit boosting (**Figure 7E**). However, we did not observe persistence of any transferred T cells by day 12 post-vaccination. We observed transient tumour regressions in all mice that received CAR-T cells, but there was no survival benefit observed in mice receiving T cells that co-expressed BCMACAR(28ζ) and mIL-15 (**Figure 7E, F, Figure S12**). Therefore, inclusion of a native murine IL-15 transgene alongside the boosting CAR is not sufficient to enhance *in vivo* T cell persistence or anti-tumour function following VSV boosting.

## Discussion

Here, we demonstrated that a CAR targeting a non-tumour antigen can be engineered into tumour-specific T cells and used to drive T cell expansion in a tumour antigen agnostic fashion. Importantly, function through the endogenous TCR was maintained in the CAR-T cells confirming the utility of this approach. The vaccine candidate we employed, recombinant VSV, is a useful tool for stimulating the T cells via the CAR but the magnitude of the T cell expansion is limited by interferon-mediated suppression of transgene expression, presumably related to suppression of viral replication. Notably, despite robust T cell expansion following CAR-mediated boosting, the engineered T cells displayed rapid contraction. In contrast, the same T cells boosted through their TCR displayed improved persistence. Thus, the mechanism by which the T cell is activated has a marked impact on its persistence and, ultimately, anti-tumour activity. The inclusion of a survival factor in the engineered T cell product, IL-15, did not impact persistence. Thus, future studies should consider alternate CAR designs and/or alternate vaccine vectors. Importantly, in the absence of boosting, VSV vaccination appeared to drive attrition of the transferred T cells, suggesting that there is a threshold above which CAR-antigen engagement leads to robust expansion, but below which prevents engraftment of the transferred cells altogether.

Type I IFN was shown by Evgin et. al. to drive attrition of CAR-T cells via a T cell-intrinsic mechanism during combination therapy in syngeneic tumour models (42). In this study, a recombinant VSV engineered to express IFNβ caused attrition of CAR-engineered T cells through interferon-induced upregulation of CAR expression that, in turn, promoted antigen-independent signaling and upregulation of coinhibitory receptors resulting in loss of function. While our results also point to type I IFN as a limiting factor in the CAR-mediated boosting, unlike Evgin’s report, the effect of IFN in this circumstance was T cell-extrinsic and the mechanism of IFN-mediated suppression of T cell expansion was a result of IFN-mediated suppression of transgene expression (as noted above). Our data do not suggest that IFN rendered the CAR-T cells dysfunctional as the T cells retain the ability to produce cytokine and cause tumour regression in the absence of IFN blockade. Even though IFNAR1 blockade enabled prolonged and enhanced expression of the boosting antigen by the VSV vaccine, thus overcoming the apparent threshold for CAR-T cell activation, there was no enhancement to CAR-T cell persistence and tumour relapse occurred in most animals. Collectively, these findings underscore the complex interplay between type I IFN signaling and CAR-T cell function, highlighting the potential of transient IFNAR1 blockade to improve initial CAR-T cell responses in the context of VSV booster vaccination strategies while also pointing to the need for additional strategies to enhance T cell persistence and durable tumour control.

Previous work by our group demonstrated that endogenous T cells were required to prevent tumour relapse following TCR-mediated boosting of transferred tumour-specific T cells (16). Here, in the DUC18/CMS-5 model, transferred tERK-specific T cells dominated the T cell population 5 days following VSV-tERK vaccination, but endogenous tERK-specific T cells were more abundant at day 12 post-vaccination. In the context of the B16.F10-gp33 tumour model, adoptive transfer of P14 T cells followed by boosting with VSV-gp33 results in transient tumour regression followed by relapse, but this relapse is driven by antigen loss (48). Interestingly, although we observed similar tumour relapse kinetics following CAR- and TCR-mediated boosting in this model, relapsed tumours consistently maintained expression of gp33 antigen when P14-BCMACAR T cells were boosted with VSV-BCMA. Neither reducing the level of CAR expression on the surface of the T cells nor reduction of CAR signal strength led to improved T cell persistence and again pointed to a threshold for CAR-mediated boosting that must be met by sufficient expression of both the CAR and the boosting antigen *in vivo*. Further, persistence of the transferred T cells was only achieved following TCR-mediated boosting, leading us to consider the context in which the CAR-T cells are experiencing their boosting antigen and how this may differ from the context of TCR-driven T cell boosting, particularly in terms of the role of antigen presentation in the context of professional APCs in secondary lymphoid organs. The expression of a CAR, regardless of whether it contained ITAMs or not, also limited the persistence of the T cells. This could be due to the development of immunity against the CAR or the CD28 signaling. Future studies will investigate these possibilities.

According to our previously described mechanism of TCR-antigen engagement following vaccination with VSV expressing a peptide antigen (49), follicular B cells are primarily infected, but T cell engagement and expansion is dependent on cross-presented antigen on splenic dendritic cells. In a CAR-mediated boost, which relies on direct expression of vaccine antigen on the surface of infected cells rather than cross-presentation of peptide antigen, it is unclear whether the transferred T cells experience activation within the same immunological context. TCR-pMHC engagement on professional antigen presenting cells, such as CD11c+ dendritic cells, may involve cross-talk between endogenous virus-reactive CD4+ T cells leading to enhanced costimulation and cytokine signaling during CD8+ T cell activation. Here, in the absence of CD4+ CAR-T cells, CAR-antigen engagement may occur in an immunological milieu that is spatially or temporally separated from CD4+ T cell help, leading to limited persistence and functionality of boosted CD8+ T cells. Supporting this hypothesis is a phenomenon described as the “helpless” CD8+ T cell (50–52). According to this, CD8+ T cells that are primed in the absence of CD4+ T cell help (“helpless” CD8+ T cells) are less functional, less able to undergo subsequent clonal expansion upon restimulation, and are more likely to become exhausted or undergo activation-induced cell death. The mechanism of CD4+ help during CD8+ T cell activation has been linked to concurrent DC licensing and subsequent production of IL-12 and IL-15, which act to promote CD8+ T cell polyfunctionality, memory formation, and persistence (53). It has been reported that the “helpless” CD8+ T cell phenotype can be rescued with IL-15 (STAT5) signaling (54). We, therefore, assessed whether persistence of the boosted CD8+ CAR-T cells could be enhanced by supplementing the CAR-T cells with additional STAT5/IL-15 signaling capabilities. However, co-expression of a functional IL-15 transgene alongside the boosting CAR did not improve *in vivo* T cell persistence following VSV-BCMA boosting. Therefore, IL-15 alone may not be sufficient to overcome the need for CD4+ T cell help or costimulation by a licensed DC during boosting.

In addition to the boosting vaccine designed to express surface-expressed hBCMA antigen, we evaluated a secreted chimeric boosting antigen in which the extracellular domain of hBCMA was fused to the murine IgG1 Fc domain based on the hypothesis that this antigen could be presented on APC populations bearing the appropriate Fc receptors. However, we found that this secreted antigen was incapable of boosting CAR-T cells *in vivo* in the presence or absence of IFNAR1 blockade, despite exceptionally high serum concentrations of hBCMA achieved with the latter. This suggests that presence of the Fc-fusion boosting antigen in the serum is not sufficient for effective CAR-T cell engagement; rather, the antigen must be stably associated with target cell membranes. Others have shown that CARs can be engaged with a soluble antigen, but only when the antigen is capable of mediating CAR dimerization (55). Therefore, to facilitate T cell activation with a soluble boosting antigen, the ligand portion of the chimeric antigen may need to be designed as a dimer or oligomer. The soluble antigen design may also be limited by the choice of Fc domain. The IgG1 Fc domain was selected due to constitutive expression of FcγRIII and FcγRIIb on splenic APC populations and low rates of receptor-mediated endocytosis upon ligand binding (56–58). However, the low-affinity Fc-FcR interaction may prevent the stable cell membrane association required for CAR engagement and effective T cell activation.

In pre-clinical studies, 4-1BB-based second-generation CARs display enhanced T cell persistence and reduced functional exhaustion over their CD28-based counterparts (59). In the context of the VSV-mediated boost, CAR(28ζ)-T cells generally showed a more robust initial expansion following VSV boosting compared to CAR(BBζ)-T cells and we did not observe a meaningful functional benefit to one costimulatory domain over the other in terms of *in vivo* persistence or overall antitumour efficacy of boosted T cells. This may be attributed to the transient nature of boosting antigen expression in our system. It is possible that 4-1BB-based CARs provide a persistence advantage in scenarios where target antigen presence is prolonged. This distinction underscores a fundamental difference between the typical use of CARs—redirecting polyclonal T cells towards a tumour target—and employing CARs as boosting receptors, where transient CAR stimulation is leveraged to induce T cell expansion and differentiation. In other instances where CAR-T cells were boosted through a vaccine directed at the CAR (60), the vaccine antigen was a tumour-associated antigen rather than a tumour-irrelevant antigen. Thus, the antigen persisted as long as the tumour was present. Similarly, a recent report employed a CD19-targeted CAR as a boosting receptor on tumour-reactive T cells, whereby endogenous B cells expressing CD19 act as the boosting target (61). Again, the antigen will persist until the B cells are fully ablated. Thus, this work ventures into relatively unexplored aspects of CAR biology in terms of the functional consequences of short-lived engagement with a vaccine-delivered antigen.

It is also important to recognize that in our system, when we employ VSV-gp33 to engage transferred T cells through their gp33-specific TCR, we are simultaneously engaging endogenous gp33-reactive T cells that arise when the B16.F10-gp33 tumours are implanted into immunocompetent recipients. In contrast, vaccination with an irrelevant antigen, such as hBCMA, leads to engagement of only transferred BCMACAR-T cells. In this way, comparing CAR- and TCR-mediated boosting may be confounded by the role of endogenous antitumour T cells in suppressing gp33 antigen expression in relapsed tumours. Further, if endogenous gp33-specific T cells are critical in sustaining durable remission of gp33-expressing tumour cells, then a truly “universal” or antigen-agnostic booster vaccine design may not be an effective approach, as we do not observe expansion of endogenous gp33-specific T cells in the context of CAR-mediated boosting. Therefore, a boosting strategy that engages both the transferred T cells and the endogenous tumour-reactive T cells may be required to mediate durable tumour regressions, at least in the absence of transferred T cell persistence.

In summary, we have demonstrated that tumour-specific T cells can be effectively boosted using a recombinant OV vaccine that engages a CAR, driving robust expansion of transferred cells in lymphoreplete animals. This CAR-mediated T cell expansion leads to transient regressions of aggressive murine tumours, highlighting its potential in enhancing adoptive cell therapy in the treatment of solid tumours. However, it is clear that the long-term functional outcomes of TCR-mediated boosting are not fully recapitulated with our paired universal boosting CAR/vaccine system. This underscores the need for a deeper understanding of the biology underlying CAR-mediated boosting, how this differs from TCR-mediated boosting, and the role of endogenous T cells simultaneously engaged by the booster vaccine. Such insights will be crucial for informing the design and application of an off-the-shelf synthetic boosting receptor and boosting vaccine pair.

## Acknowledgements

All figures and supplementary figures were created in BioRender (https://BioRender.com) with an academic license. We also thank A.J. Robert McGray for helpful discussions related to murine T cell manufacturing.

## Author contributions

Conception and design of study: RB and JLB. Acquisition of data: RB, CGM, MI, DC, and CLB. Analysis and interpretation of data: RB, CGM, MI, DC, CLB, CMS, SRW, JAH, and JLB. Provision of protocols and materials: RM, NK, JCB, BDL, SRW, and YW. Drafting and revising the manuscript: RB and JLB. All authors have approved the final article.

