## Supplemental figures and tables for "Boosting of CAR-T cells with rhabdovirus is limited by type I interferon and rapid contraction"

### Supplementary figures and tables

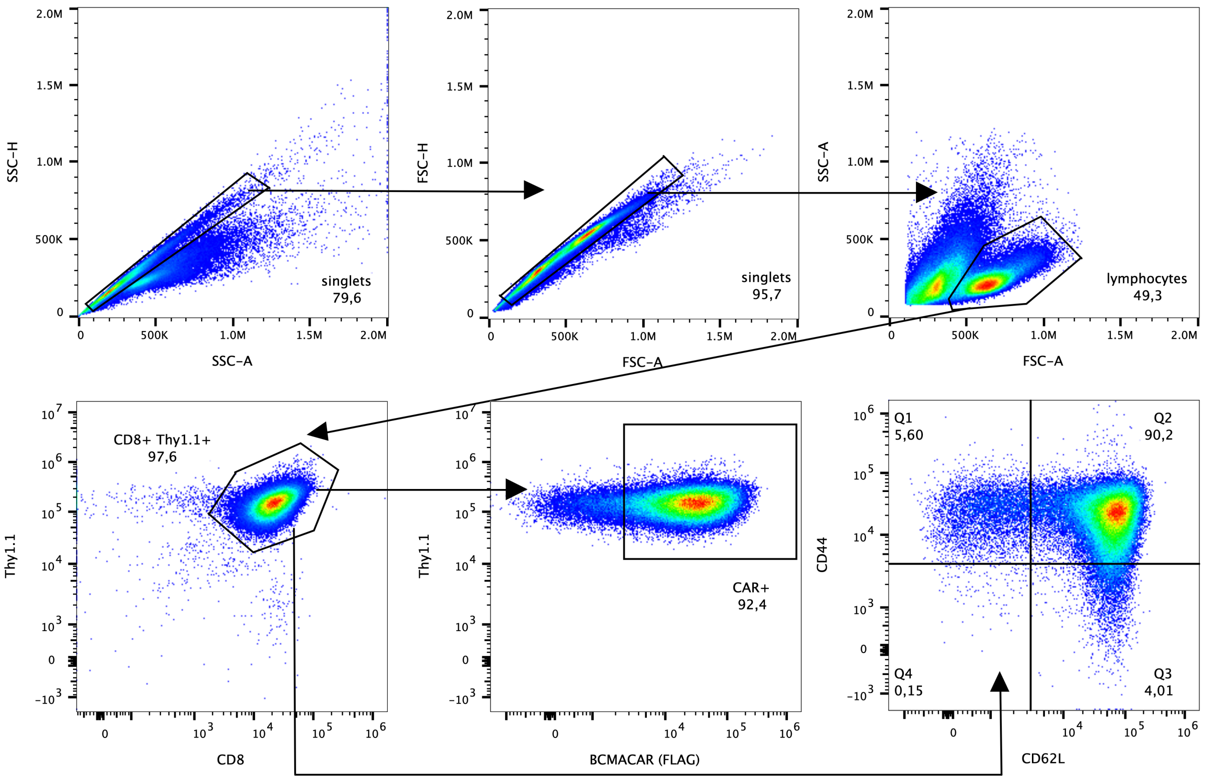


**Figure S1. Gating strategy for phenotypic analysis of murine CAR-T cells following *ex vivo* expansion.**

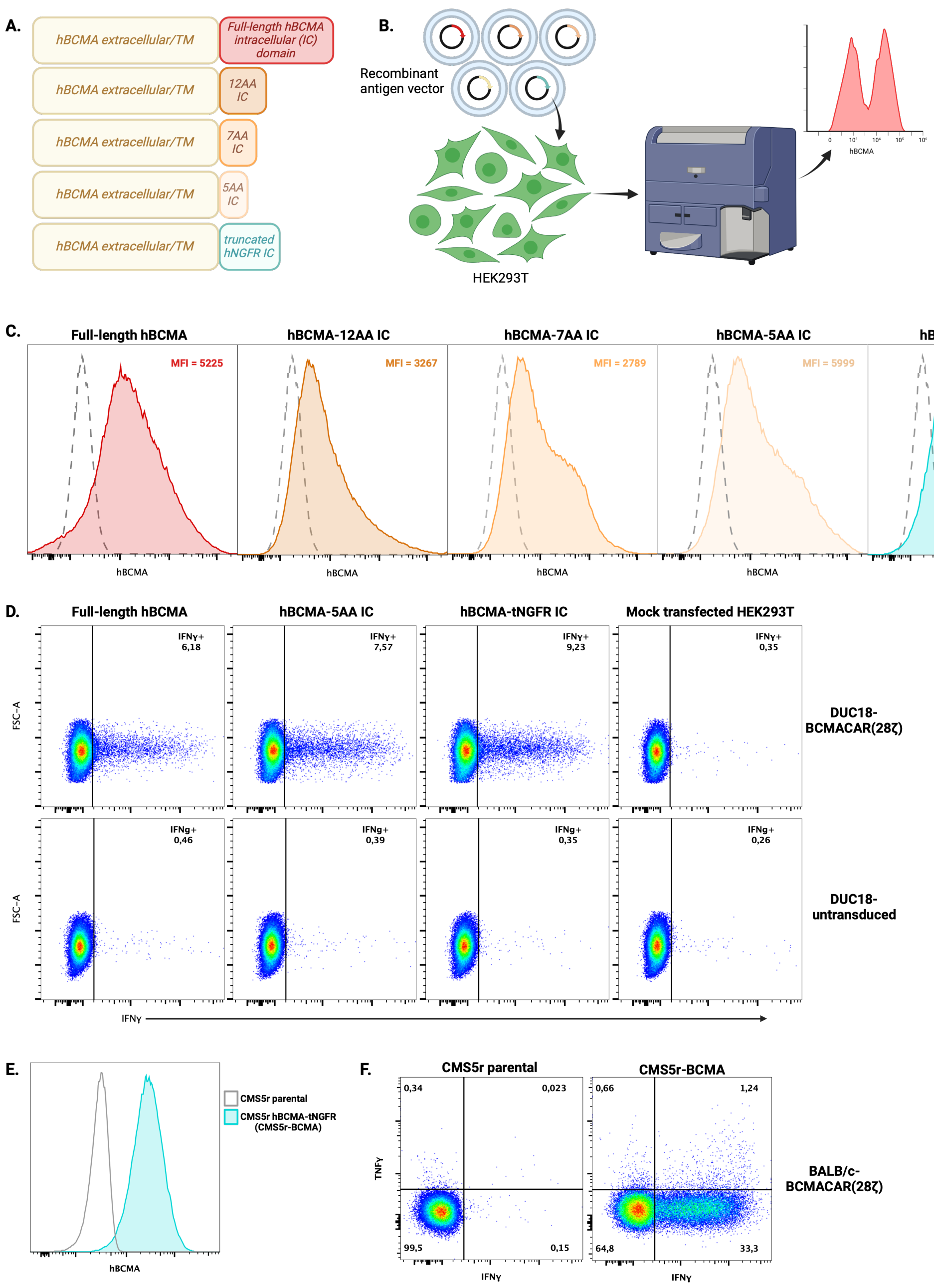


**Figure S2. Design and validation of a surface-expressed boosting antigen. A.** Schematic representation of surface-expressed human BCMA antigen designs. Different truncations were made in the intracellular domain of human BCMA to prevent signaling; expression and function of each of the variants were compared to select the antigen to go forward with. **B.** HEK293T cells were transfected with each BCMA antigen plasmid and Lipofectamine 2000. **C.** After 24 hours, cells were collected and stained with a fluorophore-conjugated anti-human BCMA antibody and antigen expression was evaluated relative to non-transfected control cells (dashed line) by flow cytometry. **D.** Cytokine production following stimulation of BCMACAR-T cells. Transfected HEK293T cells were used to stimulate BCMACAR-T cells and cytokine production was assessed by intracellular cytokine staining (ICS). T cells and target cells were co-incubated for 2 hours alone followed by 4 hours in the presence of Brefeldin A (BD GolgiPlug). Cells were collected and stained with a fixable viability dye and for surface markers (Thy1.1, CD4, CD8) and intracellular cytokines (TNFα, IFNγ) with fluorescently conjugated anti-mouse antibodies prior to flow cytometry. Events in these plots were gated down from live cells > lymphocytes > single cells > Thy1.1+ cells > CD8+. **E.** Lentivirus containing the hBCMA-NGFR transgene and a puromycin resistance gene was used to stably express the surface-expressed hBCMA-NGFR recombinant antigen in CMS5r cells. Transduced cells were selected with puromycin in culture media and grown out to establish a new engineered cell line (CMS5r-BCMA). Surface expression of hBCMA-NGFR was assessed by flow cytometry. **F.** BALB/c BCMACAR-T cells were stimulated with CMS5r parental and CMS5r-BCMA cell lines and cytokine production was assessed as in panel **D.**

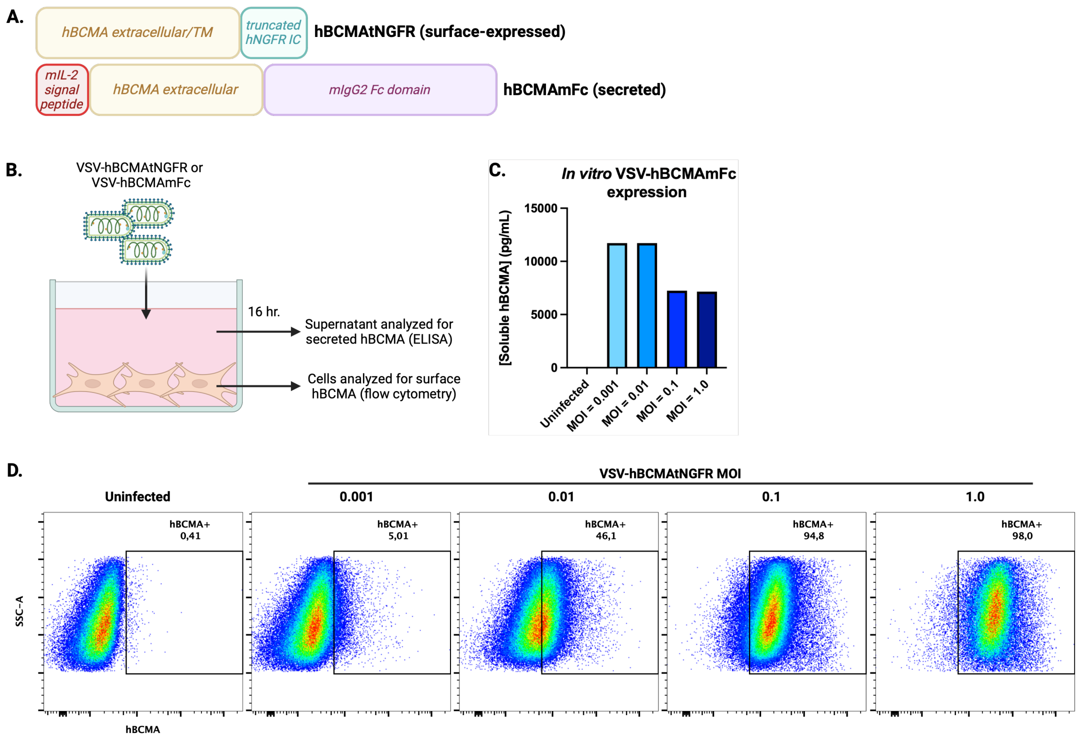


**Figure S3. Recombinant BCMA antigen expression by VSV-BCMA vaccine vectors.** **A.** Schematic representation of hBCMAtNGFR and hBCMAmFc recombinant boosting antigens. **B.** CMS5 cells were infected at various MOIs with VSV-hBCMAtNGFR and VSV-hBCMAmFc and incubated for 16 hours prior to collection of cells and supernatants, respectively, for analysis. **C.** Quantification of soluble hBCMAmFc present in supernatants of infected cells by hBCMA-ELISA (R&D Systems). **D.** Flow cytometric evaluation of hBCMAtNGFR surface expression on CMS-5 cells infected at various MOIs. Events shown were gated down from live cells by applying a fixable viability dye prior to anti-hBCMA staining

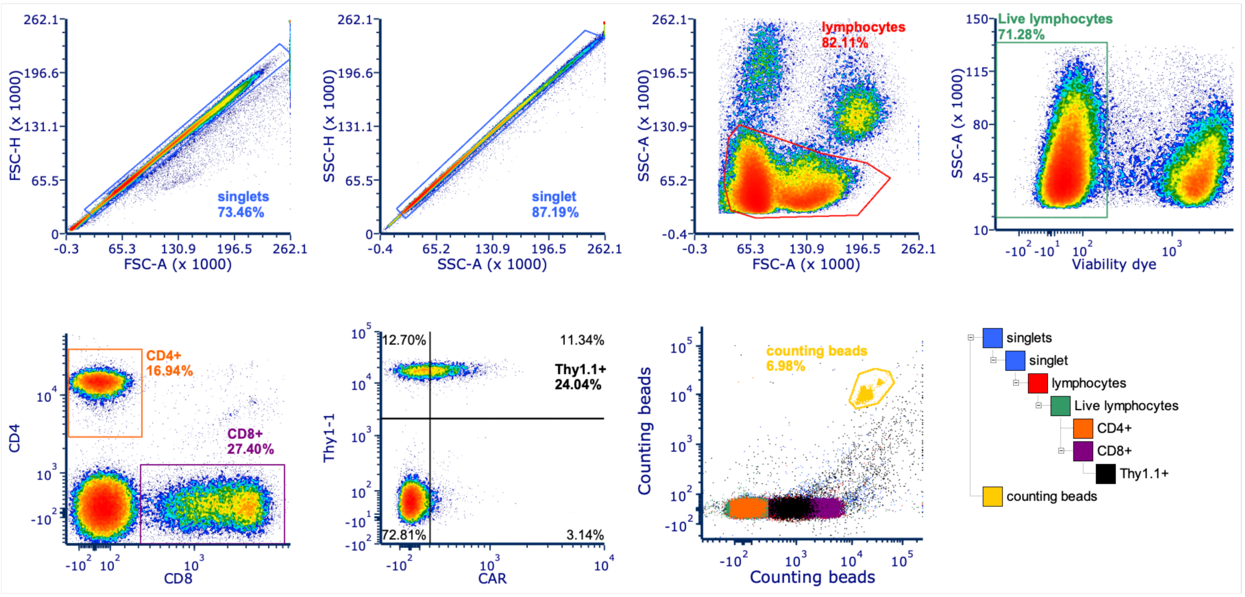


**Figure S4. Gating strategy for peripheral blood evaluation and quantification of transferred (Thy1.1+) cells by flow cytometry.**

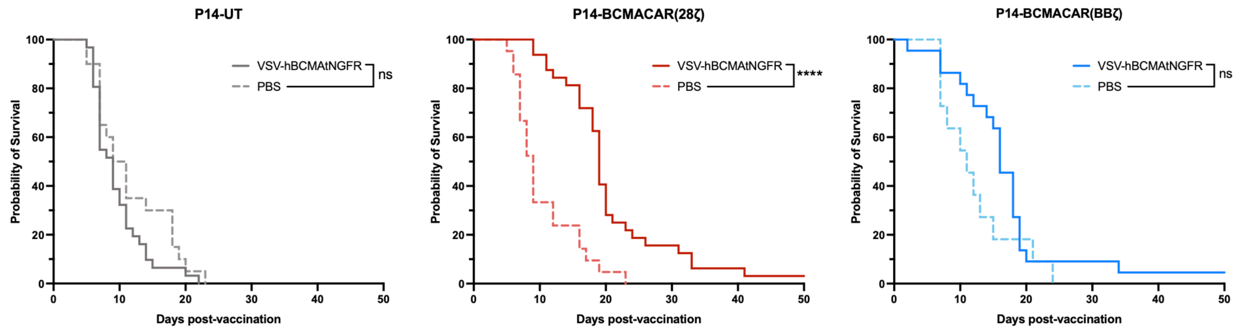


**Figure S5. Survival analysis following ACT/VSV vaccine combination therapy in the B16.F10-gp33 tumour model.** 1 x $10^6$ cryopreserved/thawed P14 CAR-T cells or P14 T cells were administered intravenously 8 days after B16.F10-gp33 tumours were implanted in C57BL/6 recipients. 24 hours after ACT, 2 x $10^8$ PFU of VSV-hBCMAtNGFR or VSV-control was administered intravenously (**Figure 2A**). Survival of tumour-bearing mice was monitored from the time of vaccination to the time of tumour ulceration or tumour volume >1000 $mm^3$.

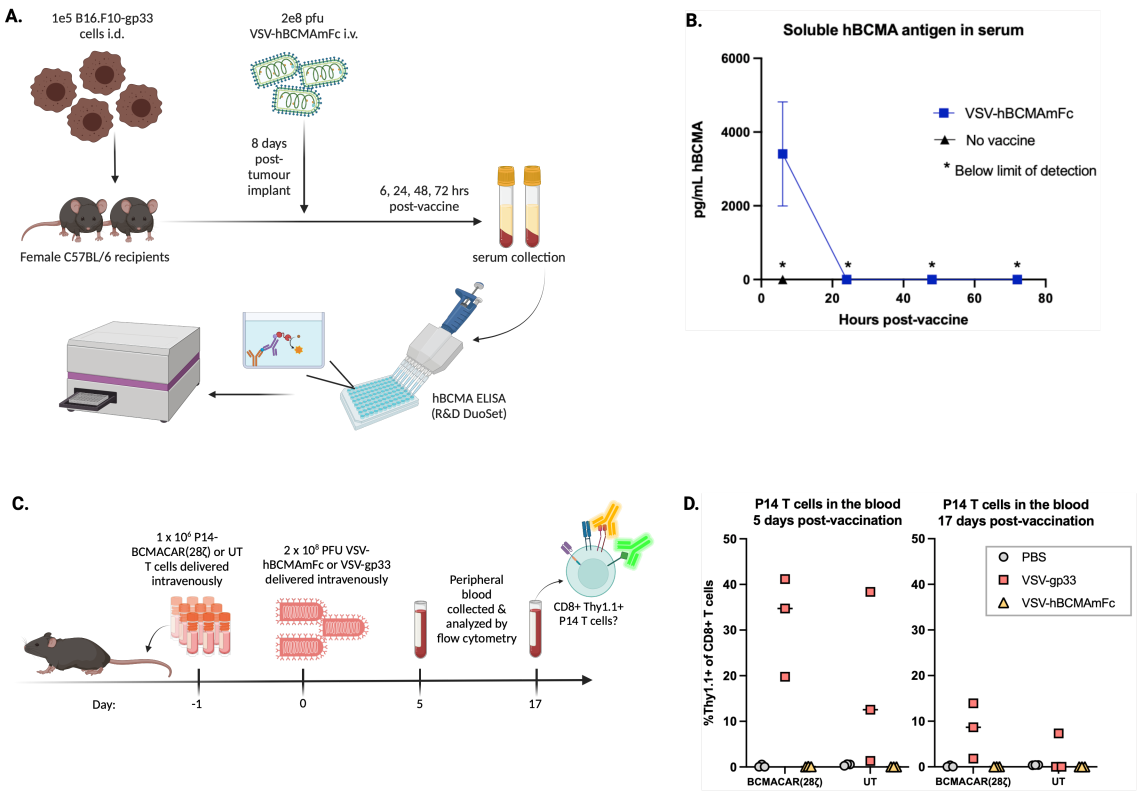


**Figure S6. *In vivo* evaluation of the VSV-hBCMAmFc soluble antigen vaccine. A.** B16.F10-gp33 tumour-bearing C57BL/6 mice were vaccinated with 2 x $10^8$ plaque-forming units of VSV-hBCMAmFc. At various times post-vaccination, serum was collected from the mice. **B.** Soluble human BCMA was quantified in the serum using a hBCMA sandwich ELISA. **C.** Schematic indicating experimental timeline and setup. **D.** 5- and 17-days post-vaccination, peripheral blood was assessed for the presence of transferred Thy1.1+ P14 cells by flow cytometry. PBMCs were stained with a fixable viability dye and fluorescently conjugated anti-mouse antibodies against surface markers prior to FACS analysis. Plots represent the percentage of total CD8+ T cells in peripheral blood that express the congenic marker, Thy1.1, in each C57BL/6 recipient post-boost (3/group).

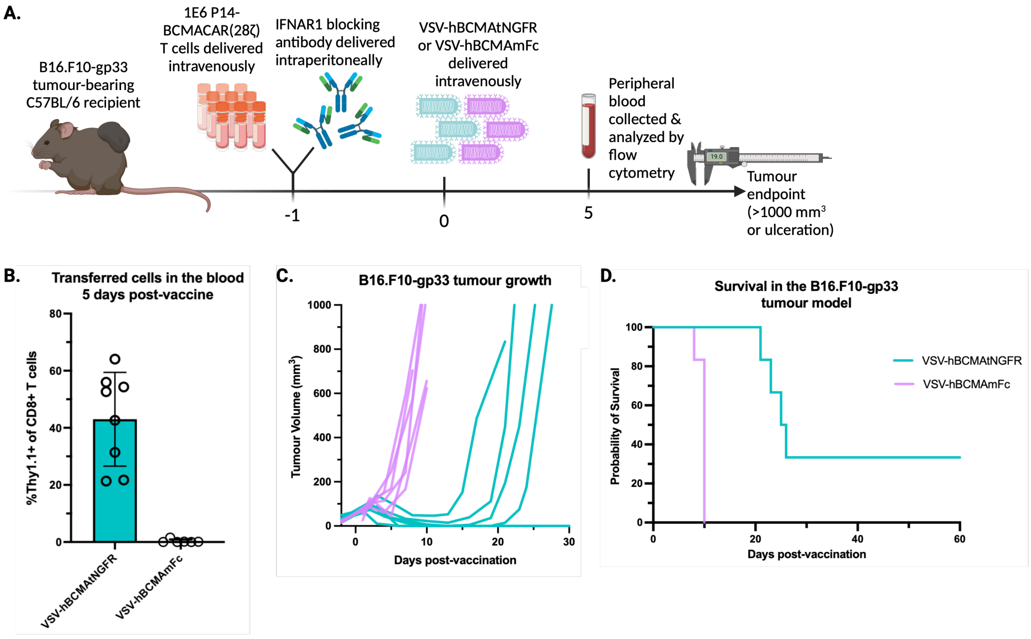


**Figure S7. VSV-hBCMAmFc still fails to boost BCMACAR-T cells in the presence of systemic IFNAR1 blockade. A.** 1 x $10^6$ P14-BCMACAR T cells were administered intravenously 8 days after B16.F10-gp33 tumours were implanted in C57BL/6 recipients. 20 hours prior to OVV, mice were given 1 mg αIFNAR1 blocking antibody. 24 hours after ACT, 2 x $10^8$ plaque-forming units of VSV-hBCMAtNGFR or VSV-hBCMAmFc was administered intravenously. **B.** 5 days after vaccination, non-terminal bleeds were collected from the facial vein of each mouse. Following ACK lysis to remove erythrocytes, cells were stained with a fixable viability dye and a surface antigen cocktail (αCD8, αCD4, αThy1.1, and αFLAG [CAR]) for quantification of cell populations by flow cytometry. **C.** Tumour size was monitored by manual caliper measurement (length x width x height) every 2-3 days for the duration of the experiment. **D.** Survival from the time that VSV vaccination was administered.

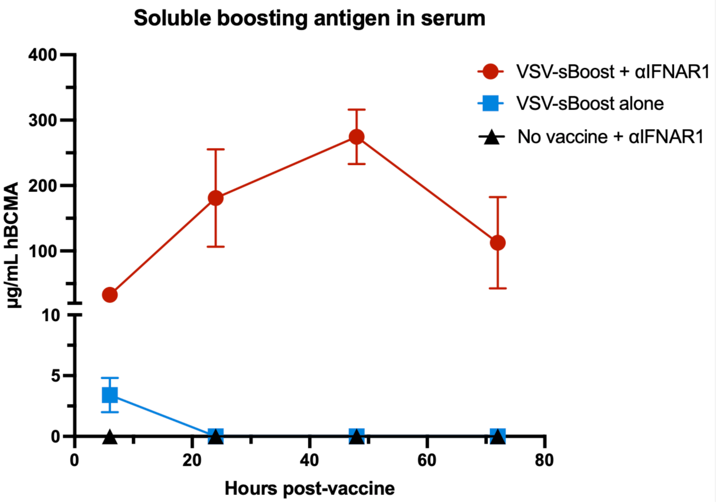


**Figure S8. VSV-hBCMAmFc antigen expression kinetics in the presence vs. absence of IFNAR1 blockade.** B16.F10-gp33 tumour-bearing C57BL/6 mice (n = 4) were administered 0.5 mg αIFNAR1 blocking antibody intraperitoneally. 20 hours later, mice were vaccinated with 2 x $10^8$ plaque-forming units of VSV-hBCMAmFc. At various times post-vaccination, serum was collected from the mice. Soluble human BCMA was quantified in the serum using a hBCMA sandwich ELISA kit. Serum samples were diluted 1:100 and run in triplicate**.**

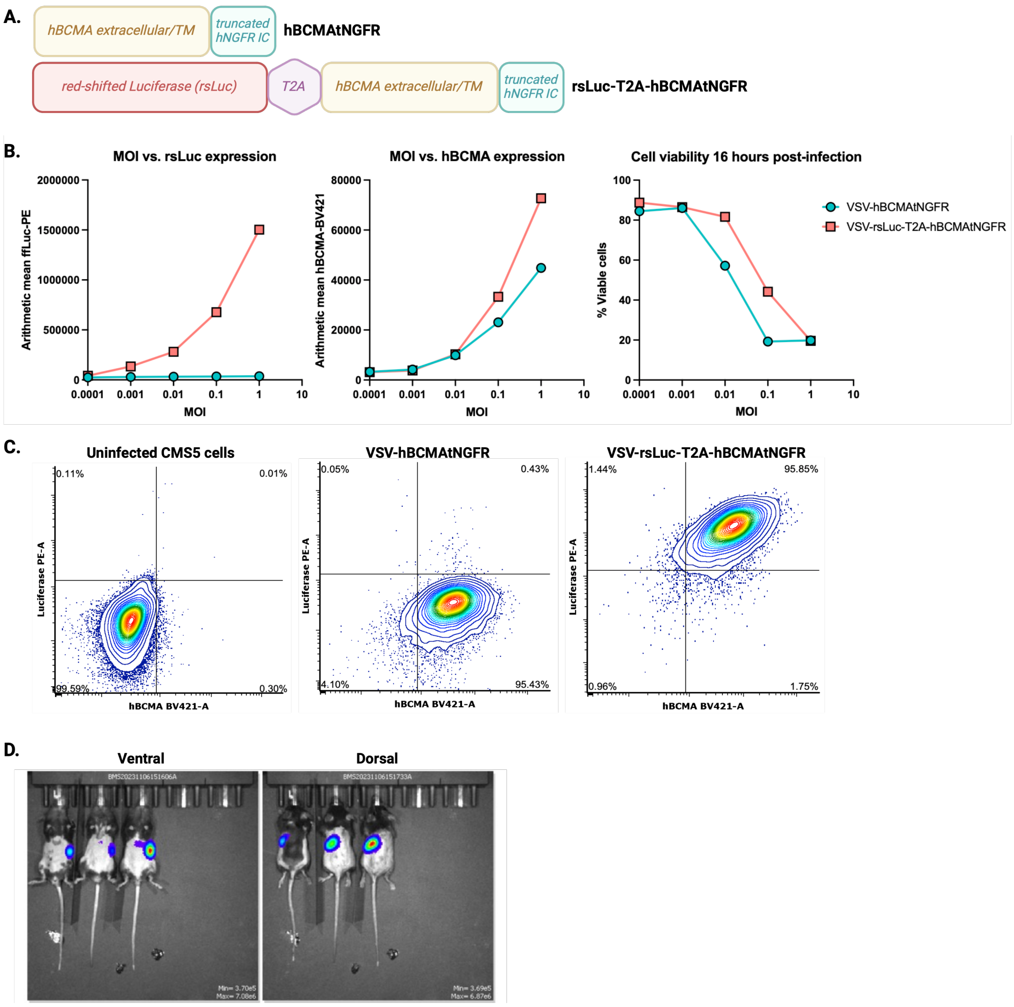


**Figure S9. Generation and validation of dual-expression hBCMAtNGFR/red-shifted luciferase (rsLuc) constructs for in vivo VSV tracking studies. A.** Schematic representation of single expression and dual-expression antigen constructs. **B.** Representative flow plots for co-expression of hBCMAtNGFR and rsLuc transgenes in CMS5 cells infected with VSV at an MOI of 1.0 for 16 hours. **C-E**. Relative expression of rsLuc, hBCMAtNGFR, and viability of infected CMS5 cells at different MOIs after 16 hours of infection, determined by flow cytometry. **F.** IVIS imaging of naïve C57BL/6 mice 6 hours post-vaccination with VSV-hBCMA/rsLuc viruses. Mice were administered αIFNAR1 (MAR1-5A3) antibody intraperitoneally 24 hours prior to vaccination.

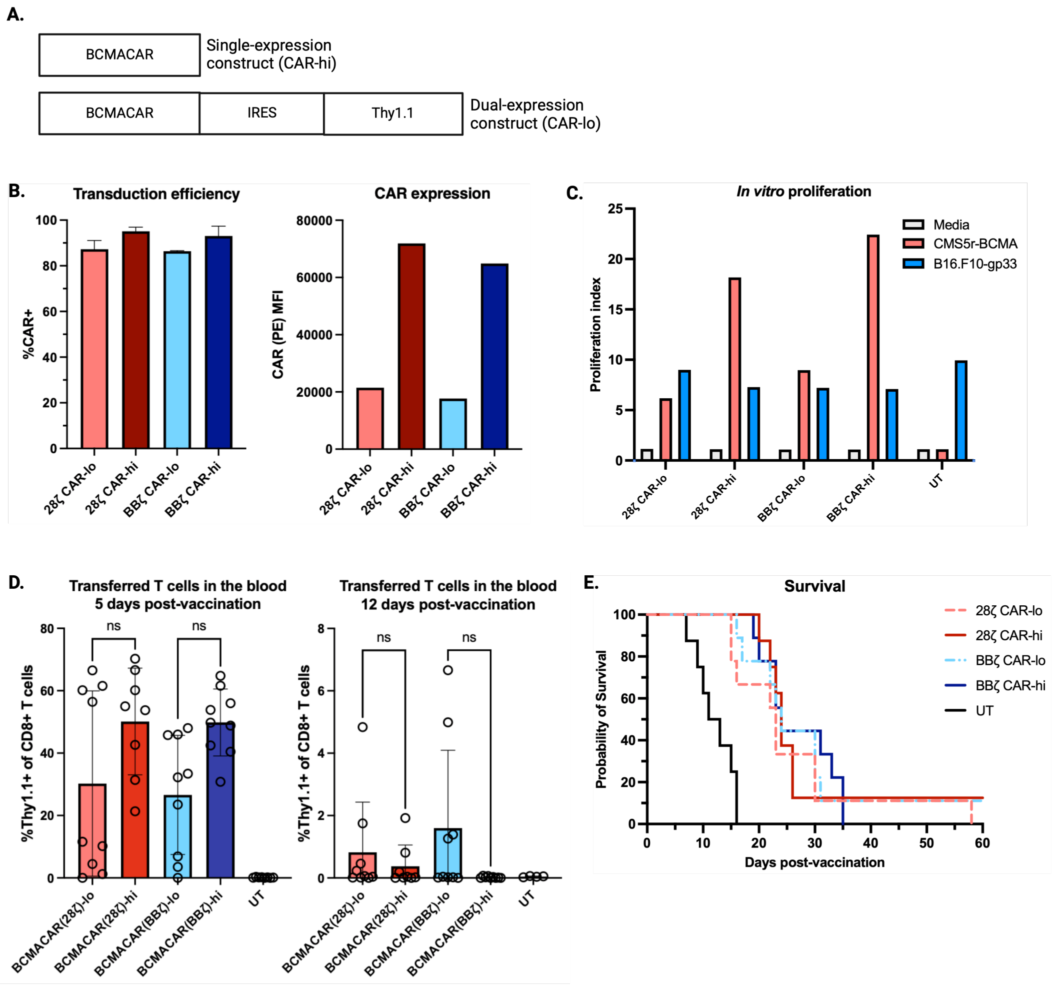


**Figure S10: Evaluating the impact of CAR expression level on P14 T cell functionality and *in vivo* persistence.** **A.** Schematic of single-expression (CARhi) and dual-expression (CARlo) construct design. **B.** P14 T cells were activated and transduced with BCMACAR constructs. On day 7 of culture, a sample of T cells from each culture was stained with fluorophore-conjugated antibodies specific for murine CD8, Thy1.1, and hBCMA-Fc/αhIgG (CAR). The stained cells were analyzed by flow cytometry for the relative abundance and distribution of each marker. **C.** CellTrace Violet dye dilution assay of P14 T cells. T cells were stained with CellTrace Violet (CTV) and co-cultured with CMS5r-BCMA or B16.F10-gp33 tumour targets for 4 days followed by flow cytometric analysis. Proliferation index of live CD8+ Thy1.1+ cells was determined by FCS Express software based on CTV dye dilution patterns. **D & E**. 1 x $10^6$ P14 CAR-T cells or UT P14 T cells were administered intravenously 8 days after B16.F10-gp33 tumours were implanted in C57BL/6 recipients. 20 hours prior to vaccination, αIFNAR1 blocking antibody was administered. 24 hours after ACT, 2 x $10^8$ plaque-forming units of VSV-BCMA was administered intravenously. **D.** 5 and 12 days after vaccination, non-terminal bleeds were collected from the facial vein of each mouse. Following ACK lysis to remove erythrocytes, cells were stained with a fixable viability dye and a surface antigen cocktail (αCD8, αCD4, αThy1.1, and αFLAG [CAR]) for quantification of cell populations by flow cytometry. **E.** Survival from the time that VSV vaccination was administered.

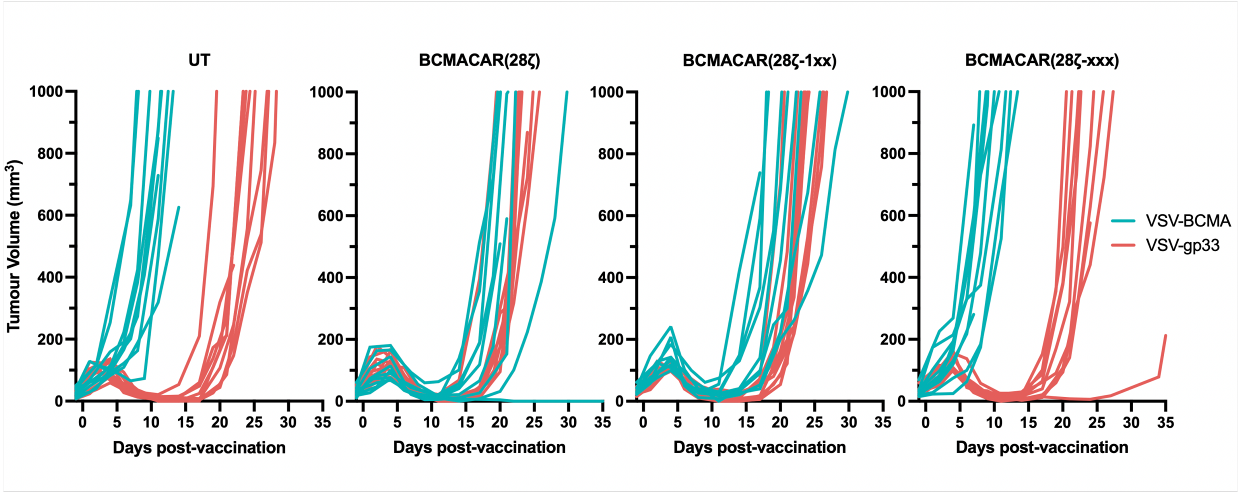


**Figure S11: Tumour growth following treatment with P14-BCMACAR ITAM-KO T cells.** 1 x $10^6$ P14 BCMACAR-T cells were administered intravenously 7 days after B16.F10-gp33 tumours were implanted in C57BL/6 recipients. 20 hours prior to vaccine, mice were given αIFNAR1 blocking antibody. 24 hours after ACT, 2 x $10^8$ plaque-forming units of VSV-hBCMAtNGFR was administered intravenously. Tumour size was monitored by manual caliper measurement (length x width x height) every 2-3 days for the duration of the experiment. Curves that truncate prior to reaching the humane endpoint of 1000 $mm^3$ are indicative of ulceration events.

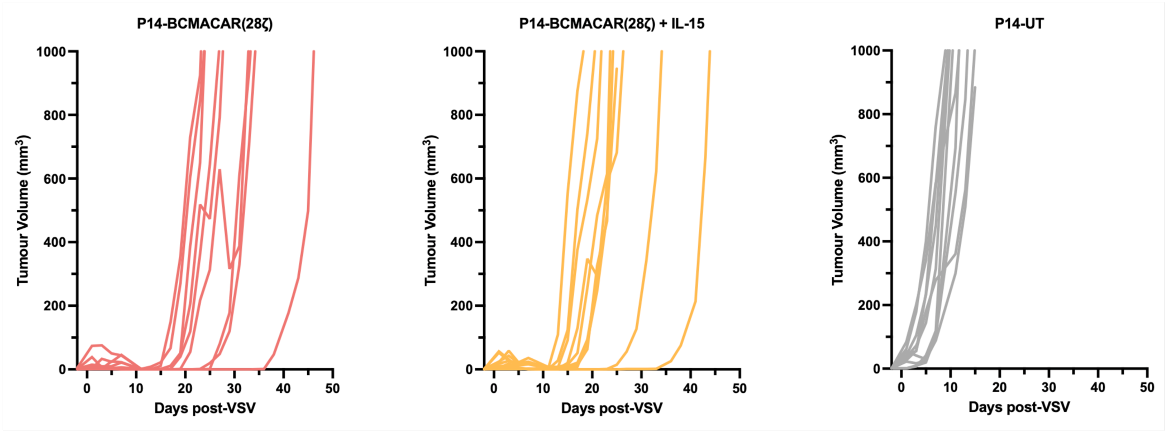


**Figure S12: Tumour growth following treatment with P14-BCMACAR/mIL-15 dual-expressing T cells.** 1 x $10^6$ P14 BCMACAR-T cells were administered intravenously 7 days after B16.F10-gp33 tumours were implanted in C57BL/6 recipients. 20 hours prior to vaccine, mice were given αIFNAR1 blocking antibody. 24 hours after ACT, 2 x $10^8$ plaque-forming units of VSV-hBCMAtNGFR was administered intravenously. Tumour size was monitored by manual caliper measurement (length x width x height) every 2-3 days for the duration of the experiment. Curves that truncate prior to reaching the humane endpoint of 1000 $mm^3$ are indicative of ulceration events.

### Supplementary tables

***Table S1. Flow cytometry antibodies used in this study.***

| Target | Fluorophore/conjugate | Clone | Supplier | Catalogue # |
|---|---|---|---|---|
| CD16/CD32 (Fc block) | - | 2.4G2 | BD | 553142 |
| CD3ε | PerCP-Cy5.5 | 145-2C11 | BD | 551163 |
| CD3ε | Brilliant violet 605 | 145-2C11 | BD | 563004 |
| CD4 | Alexa Fluor 700 | GK1.5 | eBioscience | 56-0041-82 |
| CD8α | Pacific Blue | 53-6.7 | BD | 558106 |
| CD8α | PE-CF594 | 53-6.7 | BD | 562283 |
| CD8α | Brilliant violet 421 | 53-6.7 | BD | 563898 |
| CD8α | Brilliant violet 605 | 53-6.7 | BD | 563152 |
| CD8α | Brilliant violet 711 | 53-6.7 | BD | 563046 |
| CD90.1 (Thy1.1) | FITC | HIS51 | eBioscience | 11-0900-81 |
| CD44 | APC | IM7 | BD | 559250 |
| CD44 | Brilliant violet 605 | IM7 | BD | 563058 |
| CD62L | APC-Cy7 | MEL-14 | BD | 560514 |
| FLAG (DYKDDDDK) | Brilliant violet 421 | L5 | BioLegend | 637322 |
| IFNγ | APC | XMG1.2 | BD | 554413 |
| TNFα | PerCP-Cy5.5 | MP6-XT22 | BD | 560659 |
| IL-2 | PE | JES6-5H4 | BD | 554428 |
| IFNAR-1 | APC | MAR1-5A3 | BioLegend | 127314 |
| Phosphorylated STAT1 (pY701) | PE | 4A | BD | 612564 |
| Phosphorylated STAT5 (pY694) | Alexa Fluor 647 | 47 | BD | 612599 |
| Firefly Luciferase | Unconjugated | EPR17789 | Abcam | ab185923 |
| Rabbit IgG | PE | Polyclonal | Jackson ImmunoResearch | 111-116-144 |
| Human BCMA (CD269) | APC | REA315 | Miltenyi | 130-118-975 |
| Human BCMA (CD269) | Brilliant Violet 421 | 19F2 | BioLegend | 357519 |
| Human IgG | PE | Polyclonal | Jackson ImmunoResearch | 109-115-098 |
| Human IgG | Alexa Fluor 647 | Polyclonal | Jackson ImmunoResearch | 109-605-098 |
| Human IgG | Brilliant Violet 421 | Polyclonal | Jackson ImmunoResearch | 109-675-098 |
| Mouse IgG | PE | Polyclonal | Jackson ImmunoResearch | 115-116-146 |
